# Pregnancy loss due to early developmental defects in lupus mice expressing human TLR8

**DOI:** 10.64898/2026.02.07.701591

**Authors:** Naomi I Maria, Yunwei Xia, Chirag Raparia, Ke Lin, Shani Martinez, Zhengzi Yi, Weijia Zhang, Monowar Aziz, Ping Wang, Marta Guerra, Jane Salmon, Jenny L Sones, Arnon Arazi, Paul Hoover, Anne Davidson

## Abstract

Anti-phospholipid (APL) autoantibodies confer a high risk for adverse pregnancy outcomes, especially in Systemic Lupus Erythematosus (SLE). While human TLR8 (huTLR8) has been implicated in APL antibody-mediated placental injury *in vitro*, its *in vivo* role in pregnancy is unexplored. We report a novel mouse model of pregnancy loss in SLE-prone mice expressing huTLR8. Placental analysis revealed early developmental defects starting post-implantation, including a thin junctional zone, impaired vascularization, infarcts, and inflammation. Profound immune dysregulation was evident at E8.5 including increased myeloid cells and CD8 T cells and decreased uterine natural killer (uNK) cells. RNA sequencing revealed downregulated pregnancy-specific glycoproteins, reduced uNK cell-associated genes, and an upregulated myeloid cell signature. Bone marrow chimera studies demonstrated preferential activation of huTLR8-expressing placental Ly6C^+^ monocytes. Spatial transcriptomics at E9.5 confirmed uNK cell loss, decreased *IL15* expression by both stromal and myeloid cells, and discrete inflammatory aggregates in the maternal layers containing myeloid cells and IFNγ-expressing CD8 T cells. We propose that huTLR8, likely through myeloid cell activation and cytolytic T cell recruitment, drives placental injury in the context of SLE and APL autoantibodies. This model provides a valuable platform to dissect early pathogenic events in APL-associated pregnancy loss and identify new therapeutic targets.

## Introduction

Systemic lupus erythematosus (SLE) is a multisystem, autoimmune disease affecting mainly women of child-bearing age, that causes a significant decrease in quality of life and early mortality. Genetic studies have implicated excess production of Type I interferons as a major causative pathway in SLE (1–3). Endosomal Toll-like receptors (TLRs) 7, 8 and 9 are nucleic acid sensors of the innate immune system that trigger the production of Type I interferons and other inflammatory cytokines to protect the host from infection by pathogens. However, these TLRs are also implicated in SLE pathogenesis as they can recognize endogenous nucleic acids and drive inflammatory pathways that initiate or accelerate the development of disease (4–6).

TLRs 7 and 8 are endosomal ssRNA sensors that are expressed in different cell types and recognize different RNA motifs (7). Gain of function polymorphisms of TLR7 and TLR8 predispose to SLE (8, 9) and transgenic overexpression of TLR7 or human TLR8 (huTLR8) in non-autoimmune mice causes systemic inflammatory disease (10, 11). Excess TLR7 expression accelerates SLE and induces anti-phospholipid (APL) autoantibodies in mice via activation of B cells (10, 12–16), however, the mechanistic role of TLR8 in SLE has not been well studied in mouse models because mouse TLR8 has a 5 amino acid deletion that attenuates its RNA-binding site (17).

APL antibodies are associated with adverse pregnancy outcomes in SLE patients (18). *In vitro* studies have implicated huTLR8 in mediating APL antibody-induced pregnancy loss by amplifying the release of pro-inflammatory cytokines from human placental trophoblasts or monocytes (19–21). However, to date there has been no reliable spontaneous model of APL antibody-induced placental injury, and the *in vivo* role of huTLR8 in the pathogenesis of this syndrome is unexplored. We report here the development of a novel model of spontaneous fetal loss in SLE prone huTLR8 transgenic mice that produce APL and other SLE-related autoantibodies. Our findings indicate that this is due to an early placental developmental defect characterized by an influx of maternal immune cells, decreased uterine NK cells, loss of the placental junctional zone and attenuated development of placental vasculature resulting in placental ischemia, placental inflammation, and significant fetal loss.

## Results

### Human TLR8 expression causes spontaneous pregnancy loss

HuTLR8tg mice have been previously described (11). Of four C57BL/6 founders bearing a huTLR8 BAC transgene, three developed spontaneous autoimmunity and failed to breed successfully. The fourth founder (Clone 8) did not develop spontaneous autoimmunity but is more susceptible to induced models of autoimmunity (11, 22). We bred Clone 8 mice with Sle1.Yaa mice to generate offspring with one or two copies of the huTLR8 transgene (**Figure 1A**). Male Sle1.Yaa mice express the Yaa locus on the Y chromosome that confers two copies of TLR7, and develop nephritis with early mortality. By contrast, Sle1 females expressing only one copy of TLR7 develop autoantibodies without clinical disease. The phenotype of male Sle1.huTLR8tg.Yaa mice has been previously described (23). We observed a high frequency of maternal and fetal death in female Sle1.huTLR8tg breeders, occurring in both huTLR8tg heterozygous and homozygous dams (**Figure 1B-D**). In affected pregnancies, dead or runted pups were delivered indicating stillbirth and fetal growth restriction, respectively. In some cases, maternal death resulted from dystocia due to failed delivery of non-viable pups. In addition, litter sizes of successful in Sle1.huTLR8tg pregnancies were smaller than in Sle1 controls (**Figure 1E**). Dystocia or fetal deaths did not occur in wild type Sle1 or C57BL/6.huTLR8tg females although litter sizes were smaller in C57BL/6.huTLR8tg breeders than in C57BL/6 or Sle1 controls (**Figure 1D, 1E**). Genotyping of pups from term pregnancies revealed no difference in the percentage of males and females in any of the groups (**Figure 1F**).

**Figure 1.**
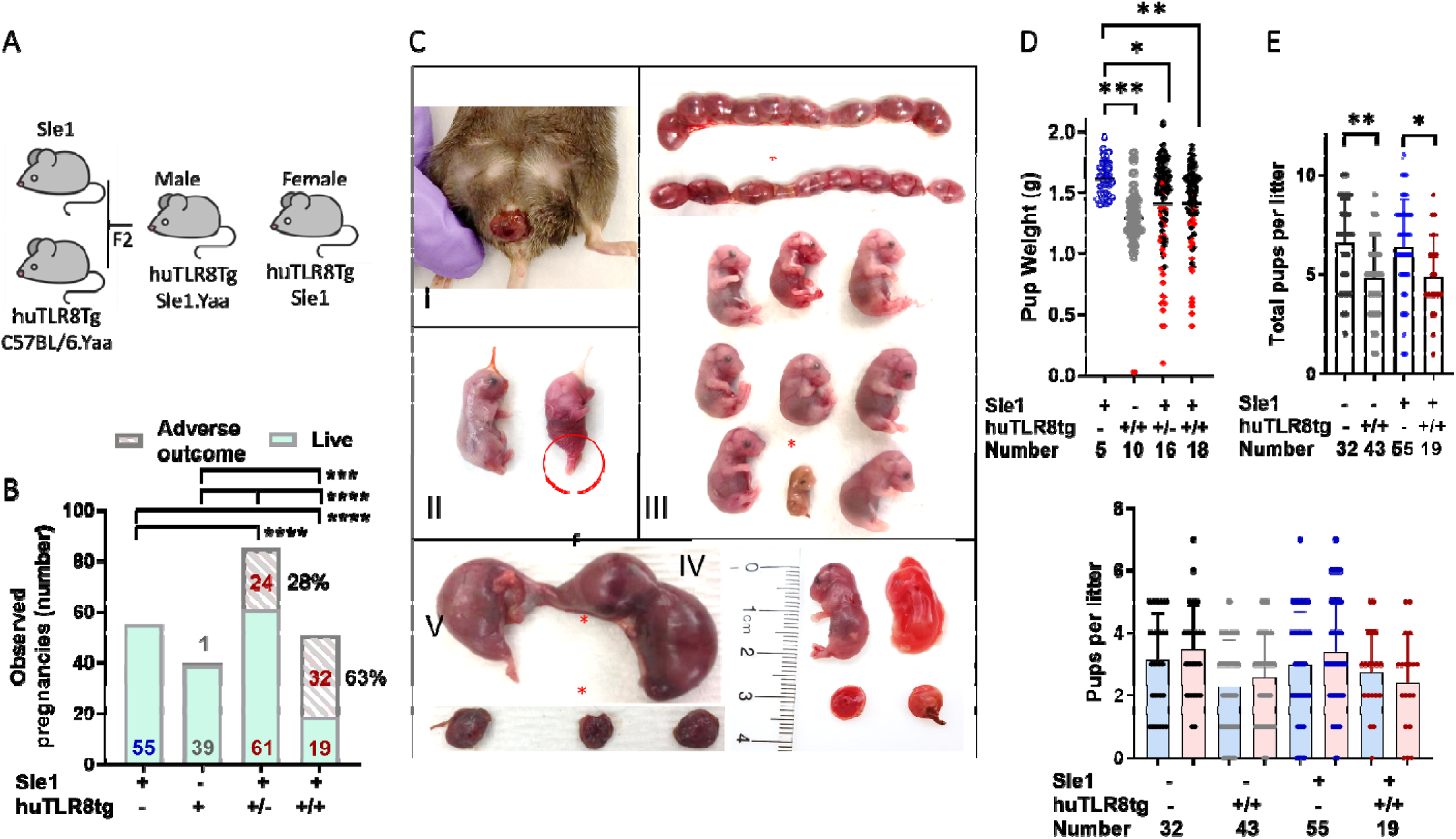
HuTLR8 induces fetal resorption and spontaneous pregnancy loss in Sle1 mice. **A)** C57Bl/6.huTLR8tg mice were crossed with Yaa males and then with Sle1 females (23). Sle1.huTLR8tg female mice express one (Sle1.huTLR8^+/-^) or two (Sle1.huTLR8^+/+^) copies of the single human TLR8 (huTLR8) transgenic locus. **B.** Outcomes of observed pregnancies. Adverse outcome was defined as maternal distress or death at delivery with evidence of an impacted pup or birth of dead or runted pups. p values calculated using Chi square analysis. **C.** I. An impacted pup in a Sle1.huTLR8^+/+^ female II. An impacted pup together with an unaffected littermate; III. Representative uteri from Sle1 (top) and Sle1.huTLR8^+/+^ (bottom) mice; IV. Growth retarded pup with an associated small placenta (*); V. A normal sized fetus and a resorbed fetus from the same pregnancy. **D.** Fetal weights (in gram) are depicted for full term Sle1 (n=5) and C57BL/6.huTLR8tg (10) control live pregnancies and Sle1.huTLR8^+/-^ (n=16) and Sle1.huTLR8^+/+^ (n=18) pregnancies with dystocia. Impacted/resorbed/dead fetuses are depicted in red. Maternal genotype and number of pregnancies are shown below the x axis. **E, F.** Total number of pups (E) and number of male (blue bars) and female (pink bars) pups (F) per live litter. Maternal genotype and number of pregnancies are shown below the x axis. Each symbol represents an individual pregnancy; bars represent the mean + SD. D-F: p values calculated using Kruskal Wallis ANOVA with Dunn’s correction for multiple analyses. *p<0.05; **p<0.01; ***p<0.001; ****p<0.0001.

### Human TLR8 transgene alters spleen phenotype and morphology in aged Sle1 mice

We evaluated the phenotype of splenic B, T and myeloid cells in females by flow cytometry at intervals from 2-11 months of age (**Supplementary Figures 1, 2**). There were no differences between C57BL/6.huTLR8tg, Sle1 or Sle1.huTLR8tg mice with respect to total cell count or B cell count (**Supplementary Figure 1A, B, Supplementary Figure 2A),** naïve B cells **Supplementary Figure 1C, 1D**), CD11b^+^ cells (**Supplementary Figure 2B, 2C**) or total CD4 and CD8 T cells (**Supplementary Figure 2D-G).** Sle1.huTLR8tg mice had a higher percentage of marginal zone (MZ) B cells than Sle1 mice but no difference in MZ cell number per spleen (**Supplementary Figure 1E, 1F**). Both Sle1 and Sle1.huTLR8tg mice accumulated germinal center B cells and plasma cells by 4-7 months (**Supplementary Figure 1G-J**) together with an increase in T follicular helper cells (T_FH_) (**Supplementary Figure 2H, 2I**). More than half of Sle1.huTLR8tg mice aged > 9 months had a decrease in germinal center B cells compared with Sle1 controls, while retaining the other B cell subsets (**Supplementary Figure 1D, 1I**). This change was confirmed by immunohistochemistry of the spleens of affected mice (**Supplementary Figure 2J**).

### Human TLR8 transgene accelerates development of autoantibody specificities in Sle1 mice

High titer APL autoantibodies have been implicated in adverse pregnancy outcomes in the context of SLE (24, 25). The presence of the huTLR8 transgene in Sle1 mice resulted in higher levels of anti-β2GP1/cardiolipin but lower levels of anti-chromatin autoantibodies at 1.5-2 months of age; these differences were no longer evident after 3-4 months of age. At all ages, Sle1.huTLR8tg mice had higher titers of autoantibodies than C57BL/6.huTLR8tg mice. Female C57BL/6.huTLR8tg mice were similar to C57BL/6 wt mice at 9 months of age (**Figures 2A-C**). No female Sle1.huTLR8tg mice developed proteinuria and kidneys from > 9-month-old mice did not display histologic evidence of glomerulonephritis or renal inflammation (**Figures 2D-G**).

**Figure 2.**
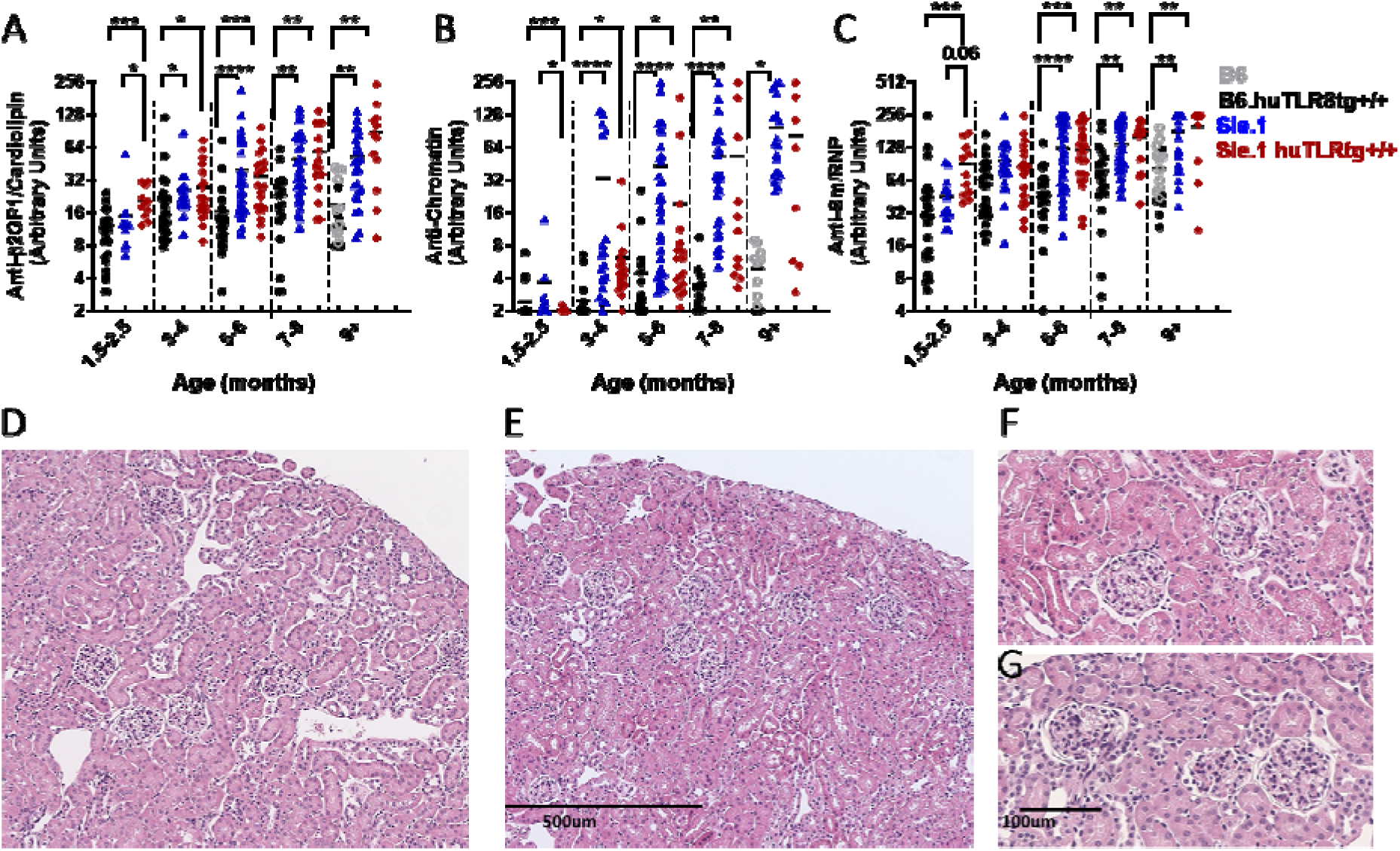
**Effect of huTLR8 on the development of autoantibodies and kidney morphology in B6.huTLR8tg, Sle1, and homozygous Sle1.huTLR8tg females**. **A-C.** Serial serum autoantibodies to β2GP1*/*cardiolipin (A), chromatin (B) and sm/RNP (C). Each symbol represents an individual mouse; horizontal lines represent the median. *P* values for each age were calculated using Kruskal Wallis ANOVA with Dunn’s correction for multiple analyses. *p<0.05; **p<0.01; ***p<0.001; ****p<0.0001. **D-G.** Representative H and E images of kidneys from 9-month-old Sle1 (D, F) and Sle1.huTLR8tg (E, G; n=5 each) show no evidence of renal injury in either strain.

### Human TLR8 expression is associated with abnormal placental morphology and inflammation

To determine the cause of fetal growth restriction and death in Sle1.huTLR8tg mice we examined term placentas of growth retarded/resorbed pups and normal sized littermates using H&E staining and immunohistochemistry. Term placentas from resorbed pups, but not littermates, showed evidence of both placental infarcts and inflammation (**Figure 3A, 3B**) with infiltration by neutrophils with a neutrophil extracellular traps (NETs)-like appearance (**Figures 3C-E**). The morphology of the affected placentas was abnormal with decreased vasculature, evidenced by loss of smooth muscle actin (SMA) staining (**Figures 3F-J**) and thickening of arterial walls in the chorionic plate as assessed by the ratio of the lumen diameter to the outer diameter of the arteries (**Figures 3K-O**). Expression of both TNF and myeloperoxidase (MPO) was significantly increased in placentas of Sle1.huTLR8.tg mice compared with controls (**Supplementary Figure 3A, 3B**) and correlated inversely with placental weight (**Supplementary Figure 3C, 3D**). There was significant thinning of the junctional zone involving both viable and non-viable pups from affected litters (**Figure 4A, 4B, 4E, 4G**). By contrast, decidual zone and labyrinth thickness were not altered in Sle.huTLR8tg placentas, with no difference in overall placental thickness (**Figures 4C, 4D, 4F)**.

**Figure 3.**
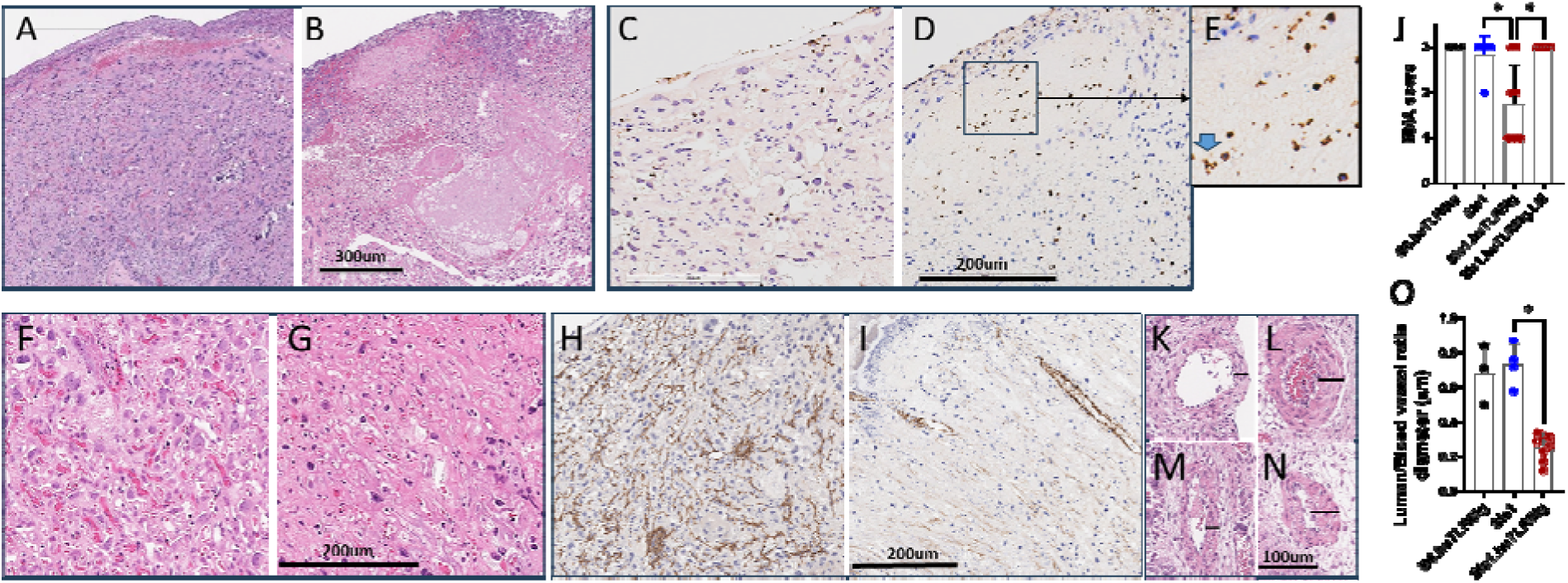
Representative morphology of placental pathologic abnormalities at term in Sle1.huTLR8tg compared with Sle1 mice. A, **B.** H&E staining shows placental infarcts and hemorrhage in the Sle1.huTLR8tg (B) compared with the Sle1 (A) placenta. **C-E.** MPO staining shows neutrophils (D) with a NETting appearance (E; blue arrow) in the Sle1.huTLR8tg placenta compared with the Sle1 (C) placenta. **F-I.** Decreased vascularity of Sle1.huTLR8tg (G, I) compared with Sle1 (F, H) placentas shown by H&E staining (F, G) and SMA staining (H, I). **J.** Semiquantitative SMA score from term placentas from the indicated strains. Resorbed/impacted Sle1.huTLR8tg placentas are shown in closed red circles and placentas from unaffected littermates (LM) are in open red circles. **K-N.** H&E evaluation of arterial wall thickness in chorionic plate in Sle1 (K, M) compared with Sle1.huTLR8tg placentas (L, N). **O.** Ratio of lumen diameter/total blood vessel diameter shows a decrease in lumen diameter in Sle1.huTLR8tg compared with control placentas. Filled circles indicate impacted pups and open circles indicate littermates. Each symbol represents the average of 2-3 vessels in an individual placenta. **J, O.** Bars represent median values. p values calculated using Kruskal Wallis ANOVA with Dunn’s correction for multiple analyses. *p<0.05.

**Figure 4.**
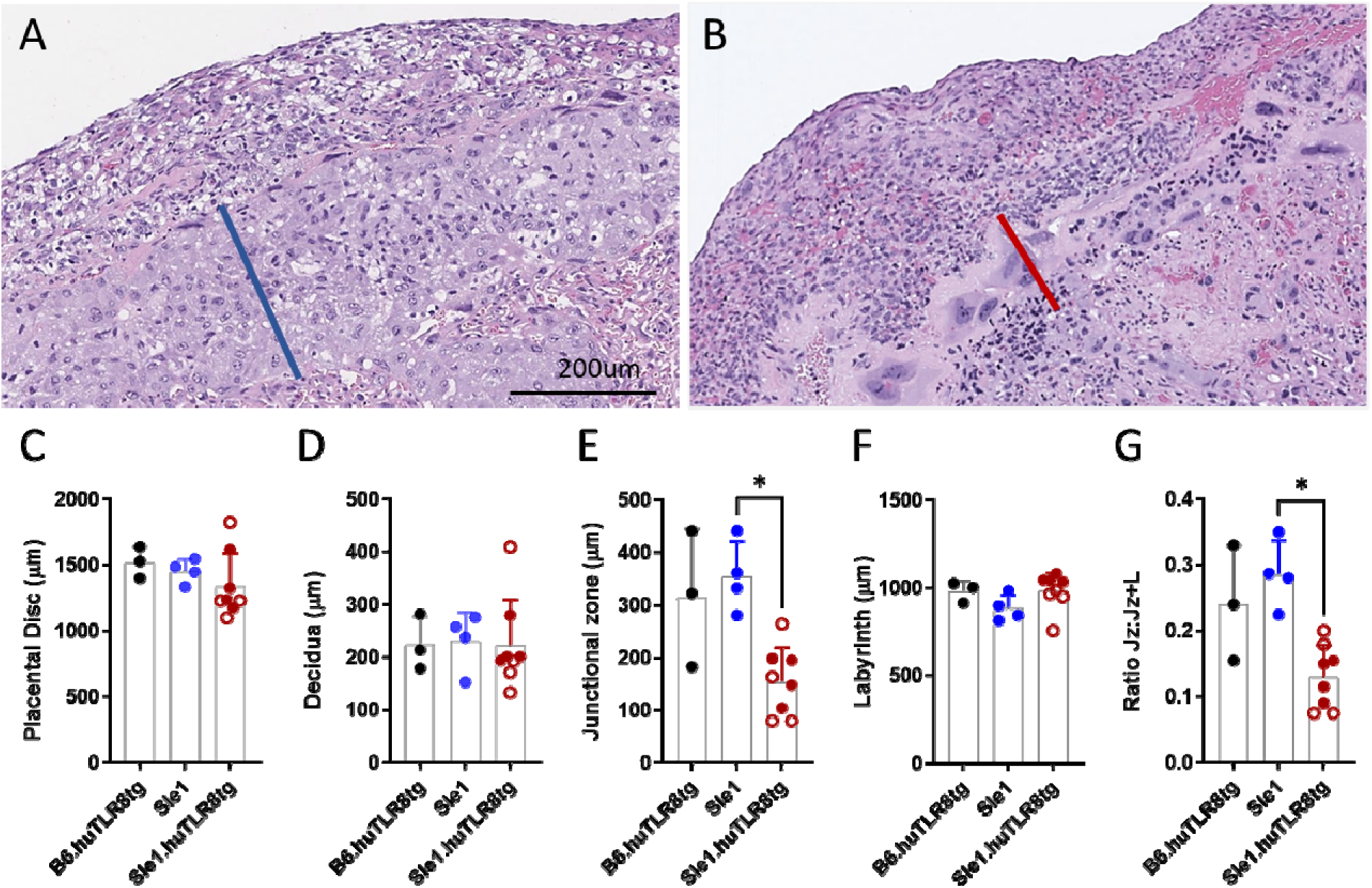
Quantification of placental layer thickness. A, **B**. Representative PAS images of placentas from Sle1(A) and Sle1.huTLR8tg (B) mice showing thickness of the junctional zone. **C-F**. Thickness of placental disc and placental layers in B6.huTLR8tg (black), Sle1 (blue) and Sle1.huTLR8tg (red) term placentas. **G.** Ratio of junctional zone to junctional zone + labyrinth thickness. Filled red circles indicate resorbed/impacted pups and open circles indicate littermates. **C-G:** Each symbol represents an individual placenta; bars represent the mean + SD. p values calculated using Kruskal Wallis ANOVA with Dunn’s correction for multiple analyses. *p<0.05.

### TNF**α** is not required for pregnancy loss in Sle1.huTLR8 mice

To assess whether TNFα was required for the observed pregnancy complications, we generated TNF deficient Sle1.huTLR8tg mice. Sle1.TNF^-/-^ mice expressing 1 copy of huTLR8 exhibited a similar frequency of dystocia and/or dead litters as Sle1.huTLR8tg^+/-^mice (8/22 = 36% vs. 24/85 = 28%) and myeloperoxidase (MPO) was present in the placentas (**Supplementary Figure 3A, 3B**). We were unable to maintain a colony of Sle1.huTLR8tg^+/+^ mice due to a high frequency of dystocia or delivery of dead litters in the females (6/8 = 75%) with runting of the surviving pups.

### Human TLR8 expression does not cause an increase in neutrophil NETs

To determine whether neutrophils from Sle1.huTLR8tg mice were prone to neutrophil NETting we stimulated BM derived neutrophils from Sle1 and Sle1.huTLR8tg mice with either PMA or the TLR8 agonist CL075. Neutrophil NETs were readily induced using both stimuli, but no difference in NETting was observed between Sle1.huTLR8tg mice and non-transgenic controls at either 2-3 or 6 months of age (**Supplementary Figure 4**)

### Human TLR8 expression is associated with placental developmental abnormalities

We next asked at which stage of pregnancy resorptions occurred in the Sle1.huTLR8tg mice. Timed breeders were harvested on E8.5, after implantation but before completion of the fetal placental unit, and on E13.5 when a fully formed placenta is present. On E8.5, the number of implanted embryos per uterus was similar in Sle1 and Sle1.huTLR8tg mice (**Figure 5A)**. However, the fetal-placental units of the transgenic mice weighed less than those of their wild type counterparts (**Figure 5B**). By E13.5, a high frequency of resorptions was evident in the transgenic mice (**Figure 5A-C**), and thinning of the junctional zone was also apparent (**Figure 5D**).

**Figure 5.**
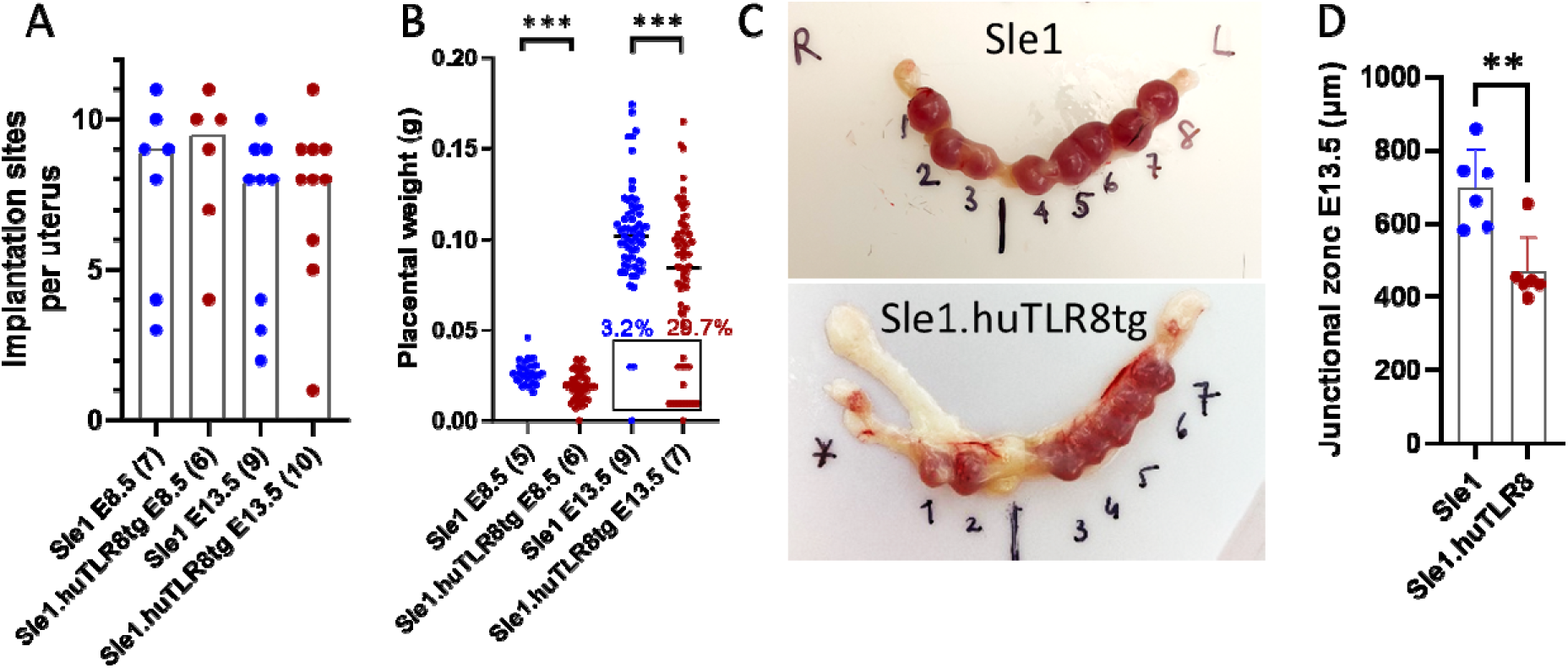
Timing of placental injury in Sle1.huTLR8tg mice. **A.** Number of implantation sites per uterus in mice of the indicated genotypes at Days E8.5 and E13.5. Bars indicate the median. **B.** Placental weights at E8.5 and E13.5 and percent resorbed placentas at Day E13.5 (box). Each symbol represents an individual placenta and number of pregnancies per group is shown on the x axis. Lines indicate median. Statistics for each age calculated using non-parametric Mann Whitney t-tests. *p<0.05, ***p<0.001. **C.** Representative uteri at Day E13.5. Star indicates a resorbed implantation site. D. Junctional zone thickness in Sle1 and Sle1.huTLR8tg placentas at Day E13.5. Each symbol represents one placenta from 6 different pregnancies in each group. Bars represent mean + SD. p value calculated using non-parametric Mann-Whitney t-test. **p<0.01.

Because preeclampsia can be associated with pregnancy loss in SLE and APS, we evaluated whole fetal-placental units from E8.5 pregnancies and isolated placentas from E13.5 pregnancies by qPCR for expression of genes known to be dysregulated in mouse preeclampsia models including Ptgs2 and Lif (involved in implantation)(26, 27), Cdkn1c (needed for trophoblast invasion)(28, 29), Csh1 and 2 (placental growth hormones)(29), VEGF, Plf and sFlt1 (involved in angiogenesis)(29–31). No differences were found in expression of these genes at either time except for a modest increase in Csh2 in Sle1.huTLR8tg fetal-placental units at E8.5 (**Supplementary Figure 5A, 5B**). Furthermore, hypertension (**Supplementary Figure 5C**) and proteinuria were not observed during the second half of monitored pregnancies from Sle1.huTLR8tg mice, and serum VEGF at E8.5-9.5 and sFlt1 levels at E18.5-19.5 were no different from control Sle1 mice (**Supplementary Figure 5D, 5E**).

To further investigate the mechanism for fetal loss in affected pregnancies, we performed bulk RNA sequencing of E8.5 fetal-placental units and E13.5 placentas of Sle1 and Sle1.huTLR8tg mice. Significant differences in placental gene expression were observed on E8.5. Major downregulated pathways in the transgenic mice involved cytolysis (Granzymes A-G and perforin), NK cell mediated immunity (Klrb1b and c and Sh2d1a, b1 and b2, Eomes, IL15, Nkp46) and cell death (Apaf1, Bcl2l2, Bcl2l11, FasL). There was also downregulation of a female pregnancy signature pathway involving members of the CEACAM family including pregnancy specific glycoproteins 16, 18, 20, 21, 23, 27 and 28 as well as CEACAMs 5 and 11-15. Major upregulated pathways included antigen presentation, inflammatory cytokines and chemokines, RNA processing, and oxidation reduction. These differences were present in both small and normal sized Sle1.huTLR8tg fetal-placental units (**Figure 6A, Supplementary Tables 1-4**). Altered expression of a subset of these genes was confirmed by qPCR (**Figure 6B-G**). At E13.5, the pregnancy signature and downregulation of prolactin family members persisted. Upregulated gene signatures at E13.5 included cell cycle and DNA replication (**Figure 6H-K, Supplementary Tables 1, 5-7**). These findings indicate a developmental defect in the placentas of Sle1.huTLR8tg mice occurring after the implantation stage.

**Figure 6.**
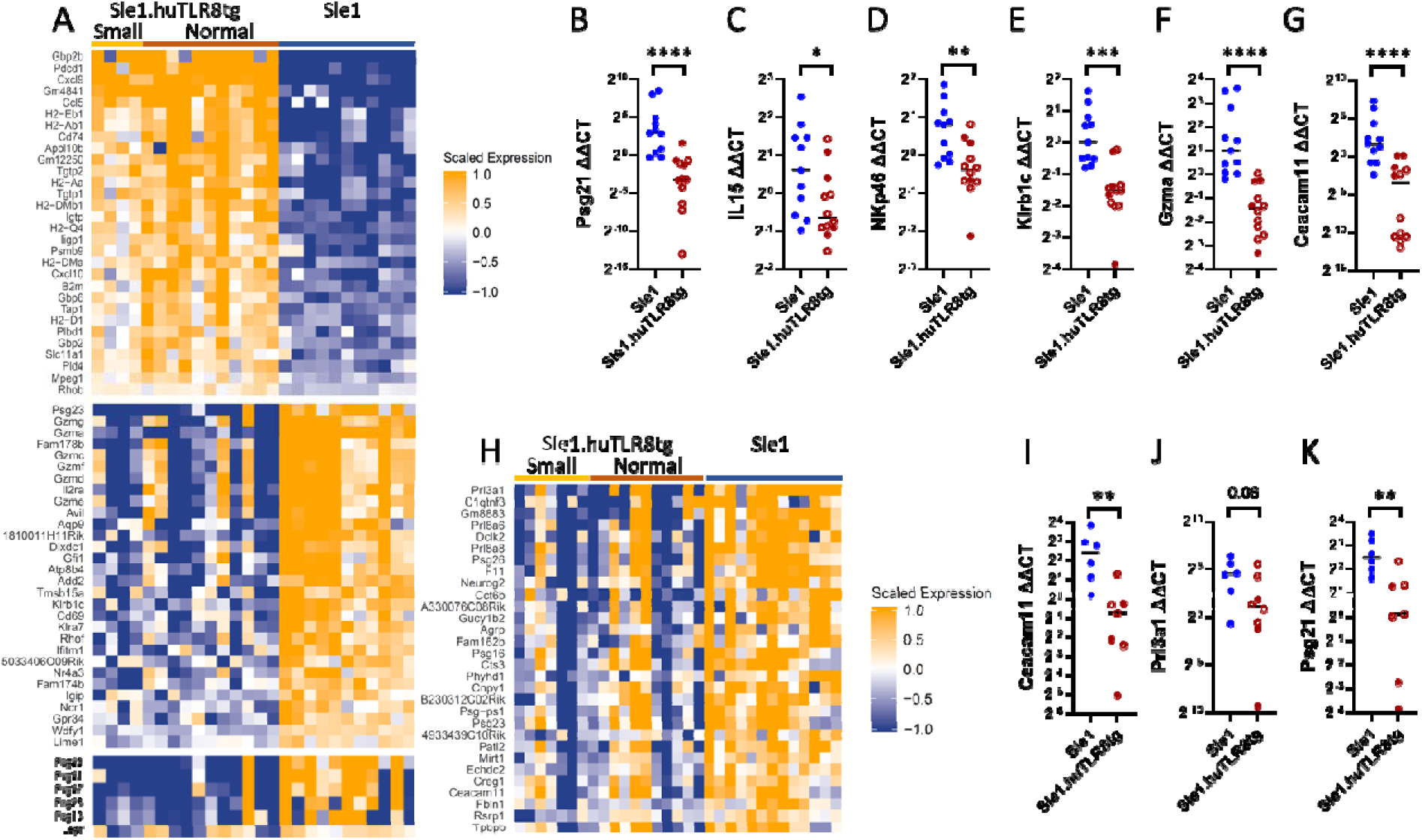
Bulk RNA sequencing of placentas. **A**. Top upregulated (upper panel) and downregulated (middle panel) genes in Sle1 and Sle1.huTLR8tg fetal-placental units at Day E8.5 show upregulation of an antigen presentation/macrophage signature and downregulation of cytotoxicity signatures. Lower panel shows downregulation of a pregnancy signature. **B-G**. qPCR confirmation of selected downregulated genes from Day E8.5 fetal-placental units. **H.** Top downregulated genes in Sle1 and Sle1.huTLR8tg placentas at Day E13.5 with persistence of the pregnancy signature. **I-K**. qPCR confirmation of selected downregulated genes from Day E13.5 placentas. B-G, I-K: Each symbol represents one placenta. Filled symbols represent small placentas. Lines indicate the median. Statistics calculated using non-parametric Mann Whitney t-tests. *p<0.05, **p<0.01, ***p<0.001, ****p<0.0001.

### Loss of the junctional zone in the presence of huTLR8 involves spongiotrophoblasts

The junctional zone separates the maternal decidua from the fetal labyrinth and contains three main cell types: a layer of trophoblast giant cells that separates the junctional zone from the decidua, glycogen cells, and spongiotrophoblasts (32). H&E staining revealed that the primary and junctional zone giant cell layers were present in the Sle1.huTLR8tg placentas (**Supplementary Figure 6A-C**) and qPCR showed no differences in expression of the giant cell specific gene placental lactogen 1 (*Pl1/Csh1*) or placental lactogen 2 (*Pl2/Csh2*) at either E8.5 or E13.5 (**Supplementary Figure 5A, 5B**). Placental glycogen production peaks at E12.5-15.5. PAS staining for placental glycogen at E13.5-14.5 showed that thinning of the junctional zone was already evident (**Figure 5D**) but placental glycogen producing cells were present (**Supplementary Figure 6C**) and RNAseq analysis of E13.5 placentas revealed no difference in expression of Gjb3 (Cx31), Gjb5 (Cx31.1), Pcdh12, Prl7b1, or Prl6a1 that are genes expressed by placental glycogen producing cells (**Supplementary Table 1**). These findings contrast with a decrease in expression of pregnancy specific glycoproteins (PSGs) that are secreted by spongiotrophoblasts, both at E8.5 and E13.5, confirmed by qPCR (**Figure 6A, 6B, 6H, 6K, Supplementary Table 1**), suggesting that the junctional zone thinning involved a loss of spongiotrophoblasts.

### Loss of uterine NK cells in Sle1.huTLR8tg placentas

Uterine natural killer (uNK) cells are a dominant immune cell type in both mouse and human placentas and are major modulators of placental vascularization and uterine spiral artery remodeling (33). Two major subsets of uNK cells are present in varying frequencies during placental development. On E8.5, most uNK cells are tissue resident CD49a^hi^ CD49b^lo^ cells whereas by E9.5-13.5, CD49a^lo^ CD49b^hi^ NK cells derive from the circulation (33, 34). Given the prominent loss of a cytolysis signature in transgenic placentas (**Figure 6A**), we visualized placental NK cells by flow cytometry and immunohistochemistry at E9.5 and E13.5. Flow cytometry demonstrated a decrease in CD4^-^CD8^-^NK1.1^+^ cells at both developmental stages in Sle1.huTLR8tg compared with Sle1 placentas (**Figure 7A, Supplementary Figure 7A)**. Using dolichos biflorus agglutinin (DBA) staining, we confirmed that there was a loss of placental DBA^+^ NK cells at E8.5 and E13.5 in Sle1.huTLR8tg compared with Sle1 placentas (**Figure 7B-F**). Using flow cytometry, we showed, after exclusion of intravascular NK cells, that there was no change in the relative representation of each uNK cell subset in the Sle1.huTLR8tg placentas (**Supplementary Figure 7B-D)** suggesting a deficiency of a required growth factor rather than a migration defect. By contrast, there was an increase in the percentage of CD8^+^ T cells in Sle1.huTLR8tg placentas compared with Sle1 controls (**Supplementary Figure 7E, 7F).**

**Figure 7.**
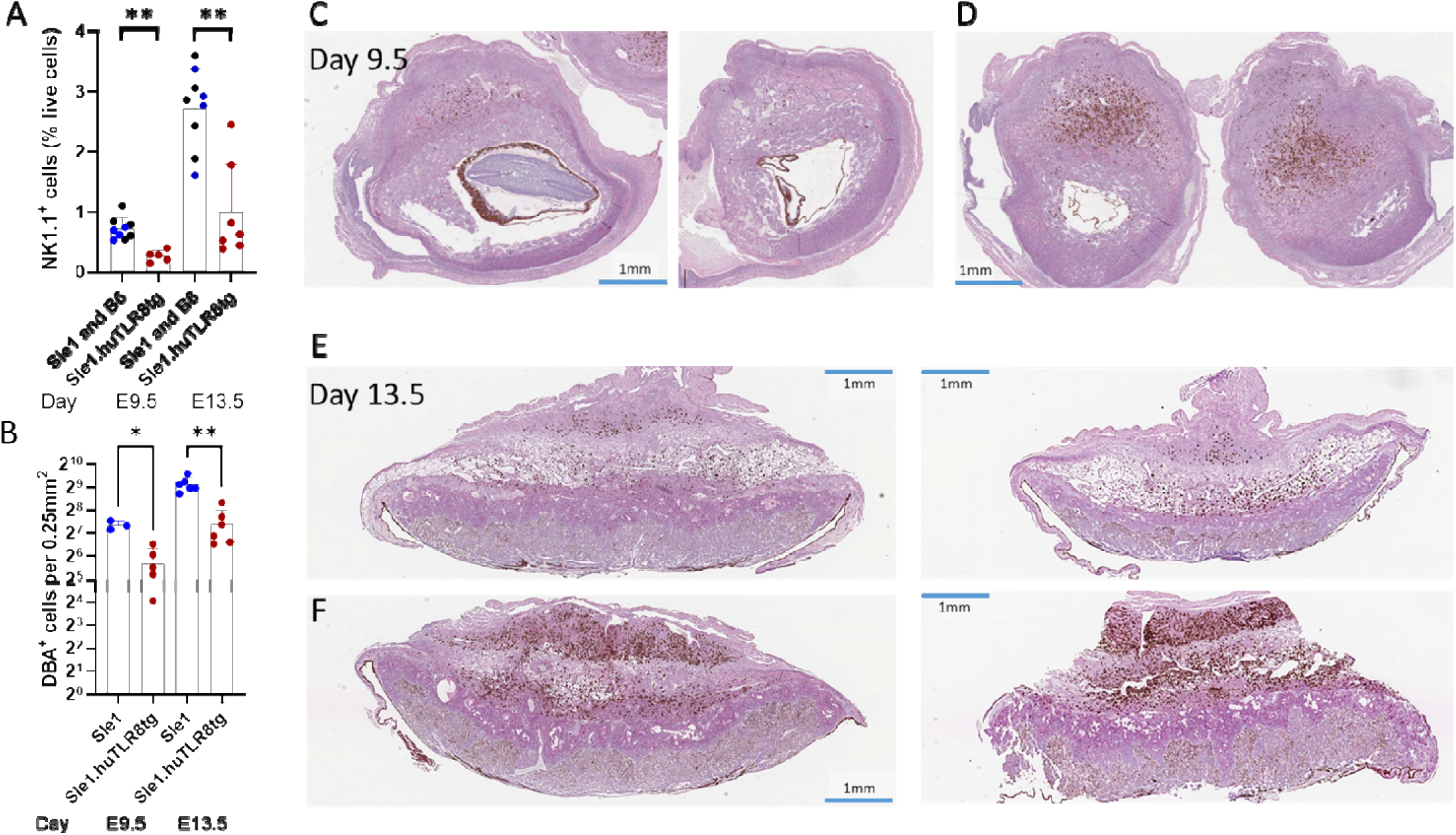
Depletion of NK cells in Sle1.huTLR8tg placentas. **A.** Percent of placental CD4^-^CD8^-^NK1.1^+^ cells in Sle1 (blue), C57BL/6 (black) and Sle1.huTLR8tg (red) mice evaluated by flow cytometry at Days E9.5 and E13.5. Each symbol indicates pooled placentas from a single uterus. **B.** Density of DBA+ placental NK cells at Days E9.5 and E13.5. Each symbol represents one placenta. A, B: Bars indicate mean + SD. Statistics for each developmental stage calculated using non-parametric Mann Whitney t-tests. *p<0.05, **p<0.01. **C-F.** Representative immunohistochemistry showing two Sle1.huTLR8tg (C, E) and two Sle1 (D, F) placentas stained with DBA at Days E9.5 and E13.5.

### Myeloid cell alterations in huTLR8tg placentas

We have previously shown that huTLR8 is expressed mainly in myeloid cells in huTLR8tg mice (23). Because we found an upregulated myeloid signature in E8.5 placentas, we also evaluated myeloid cell frequency and subset distribution by flow cytometry at E9.5, when sufficient cells were available for flow cytometry, and at E13.5 and E18.5 (**Supplementary Figure 8A**). At E9.5 the frequency of myeloid cells in the fetal-placental unit as a proportion of total live cells was higher in Sle1.huTLR8tg mice than in Sle1 controls, but this difference was no longer present at E13.5 (**Supplementary Figure 8B**). Subsetting of CD11b cells (35) at E9.5 showed no difference in distribution of the major subsets (**Supplementary Figure 8C**). By E18.5 an increase in the percentage of placental neutrophils was evident in Sle1.huTLR8tg mice compared with Sle1 mice, reflecting inflammation secondary to placental injury (**Supplementary Figure 8D**).

To determine whether there was preferential recruitment or expansion of Sle1.huTLR8tg placental macrophages we generated bone marrow chimeras using a non-irradiation method (36) by transferring congenic CD45.2 Sle1 or Sle1.huTLR8tg bone narrow cells into CD45.1 Sle1 recipients and harvested placentas at E10.5-12.5. Eight of 11 Sle1.huTLR8tg chimeric mice had at least one affected fetus compared with three of 9 control chimeras, with a higher percentage of resorbed or small pups (20.7% vs. 8.3% p = 0.03) and numerically smaller median litter size (6 vs. 9). For flow cytometry, mice were injected with anti-CD45 10 minutes prior to perfusion to exclude intravascular cells from the analysis. There was no preferential increase in representation of donor myeloid cells or any of the major myeloid subsets in the placentas, compared with blood in either Sle1.huTLR8tg or control Sle1 chimeras (**Supplementary Figure 9A, 9B**).

To explore the possibility that the myeloid cell expansion in Sle1.huTLR8tg placentas is associated with altered cell activation, we performed flow cytometry of peripheral blood and placentas of the chimeric mice using antibodies to CD86 and MHC Class II. MHCII^hi^CD11b^+^ cells were expanded in all placentas compared with blood; these cells were mostly Ly6C^+^ (classical) monocytes, and they included both circulating (CD45^+^) and tissue (CD45^-^) cells (**Supplementary Figure 9C**). Furthermore, donor (CD45.2) cells were enriched within the Ly6C^+^MHCII^hi^ but not in the Ly6C^+^MHCII^lo^ population in both blood and placentas of Sle1.huTLR8tg chimeras but no such enrichment was found in control Sle1 chimeras (**Supplementary Figure 9D, 9E**). CD86 expression was also higher in the Ly6C^+^MHCII^hi^ population of the Sle1.huTLR8tg chimeric placentas compared to the Ly6C^+^MHCII^lo^ population but there was no such difference in placentas of control Sle1 chimeras (**Supplementary Figure 9F, 9G**). Together, these data suggest that Sle1.huTLR8tg Ly6C^+^ monocytes have an activated antigen presentation phenotype.

### Spatial transcriptomics reveals localization of huTLR8 to placental myeloid cells

To determine which placental cells express the huTLR8 transgene we performed spatial transcriptomics on five Sle1.huTLR8tg and three Sle1 E9.5 fetal-placental units (**Figure 8A**) using Xenium stock primers, custom primers specific for huTLR8 and IL15, and a custom panel of additional primer sets to identify myeloid cell subsets and NK cells as previously described (35). The gene panel identified the placental layers (**Figure 8B**) as well as myeloid cells, NK cells, and lymphocytes. HuTLR8 was expressed only in the Sle1.huTLR8tg placentas (**Figure 8C)**, predominantly in CD11b^+^ myeloid cells located in both the mesometrial lymphoid aggregate of pregnancy (mLAP) and decidua (**Supplementary Figure 10A, 10B**). The spatial transcriptomic data also confirmed the loss of NK cell associated transcripts and IL15 in the Sle1.huTLR8tg placentas (**Figure 8D, 8E, Supplementary Figure 10D, 10E**). IL15 mRNA was detected both in stromal and myeloid cells in the mLAP and decidua of Sle1 mice and was decreased in stromal cells in Sle1.huTLR8tg placentas with a trend for a decrease in myeloid cells (**Supplementary Figure 10D-I**). We found a striking accumulation of immune aggregates containing CD8 T cells, some of which expressed IFNγ, together with myeloid cells and NK cells within the mLAP and, to a lesser extent, in the decidua (**Figure 8F-H**). The increase in CD8 T cells at E9.5 was confirmed by flow cytometry and persisted at E13.5 (**Supplementary Figure 7E, 7F**). These data in sum demonstrate an inflammatory phenotype in E9.5 placentas with a global decrease in IL15 expression and loss of uNK cells.

**Figure 8.**
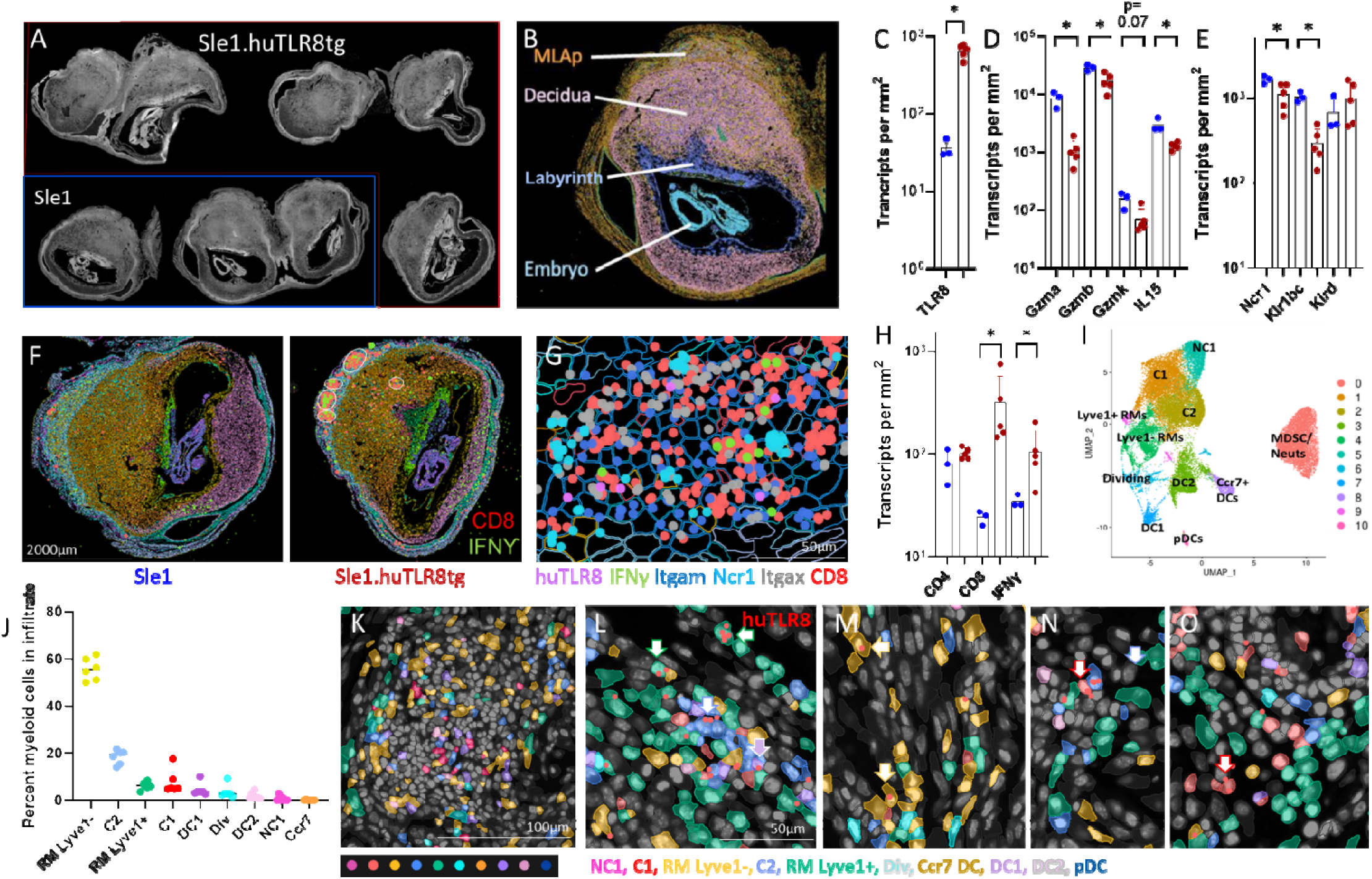
Spatial transcriptomic analysis of E9.5 placentas from Sle1 and Sle1.huTLR8tg mice. **A.** Low power image of fetal-placental units used for the experiment (DAPI staining in white). B. Placental layers identified using K means clustering. **C-E.** Enumeration of the indicated transcripts/mm^3^ in Sle1 (blue) and Sle1.huTLR8tg (red) fetal-placental units. Statistics for each gene calculated using non-parametric Mann Whitney t-tests. *p<0.05. **F, G.** Inflammatory infiltrates in the mLAP of Sle1.huTLR8tg mice contain CD8 T cells (red dots) expressing IFNγ (green dots) as well as NK cells expressing Ncr1 and myeloid cells expressing Itgam and Itgax. **H.** Enumeration of transcripts/mm^2^ in Sle1 (blue) and Sle1.huTLR8tg (red) fetal-placental units. Statistics for each gene calculated using non-parametric Mann Whitney t-tests. *p<0.05. **I.** Clustering of placental myeloid cell subsets. **J.** Percent of myeloid cells in each subset in 6 individual inflammatory infiltrates from 5 Sle1.huTLR8tg placentas. **K.** A representative inflammatory infiltrate showing the myeloid cell subsets. **L-O.** Representative images showing expression of huTLR8 (red dots) in multiple myeloid cell subsets.

To further understand which myeloid cell subsets express huTLR8, we used E9.5 scRNAseq data and our previous renal macrophage scRNAseq data to identify myeloid subclusters (**Figure 8I**) that were then mapped onto the Xenium data as previously described (35). Lyve1 positive and negative resident myeloid cells segregated in the uterine wall and mLAP respectively (**Supplementary Figure 10J, 10K**). The immune aggregates in the mLAP contained Lyve1 negative resident macrophages, as well as Ly6c positive Classical 1 and Classical 2 monocytes and cDC1 (**Figure 8J, 8K**) whereas non-classical monocytes were confined to the decidua (**Supplementary Figure 10L, 10M**). Insufficient markers were available to reliably identify neutrophils. All myeloid cell types expressed the huTLR8 transgene (**Figure 8L-O**).

### Serum levels of PSG1 in human APS and SLE pregnancies

A major gene expression signature at E8.5 and E13.5 was a loss of PSGs that are made in the junctional zone. PSGs are not detectable in the serum of pregnant mice because they are rapidly degraded (37). However, in human pregnancies, PSGs appear in the serum during the first trimester and increase up to the time of delivery. Previous studies have suggested that low levels of PSG1, the major human PSG, in the third trimester are associated with poor pregnancy outcome including low birth weight, preeclampsia or fetal death (38–40). We therefore asked whether low levels of PSG1 are associated with poor outcome in either APS or SLE pregnancies using sera obtained prospectively and longitudinally as part of the PROMISSE study (41). We found that serum levels of PSG1 were highly variable among healthy pregnancies and that there were no differences between pregnancies with either good or adverse outcome among APS or SLE patients or between healthy and APS or SLE pregnancies (**Supplementary Figure 11**).

## Discussion

SLE is associated with adverse pregnancy outcomes, particularly in the presence of anti-phospholipid antibodies (APL) (18, 42, 43). However, the underlying mechanisms for damage of the fetal-maternal unit remain poorly understood (44, 45). Previous mechanistic studies of the effector phase of APL antibody-induced placental injury used inducible or *in vitro* models (19, 46, 47). Here we identify a new spontaneous model of adverse pregnancy complications in mice with APL and other lupus-related autoantibodies, allowing study of early pathogenic events. Pregnancy loss in this model requires both *huTLR8,* with maternal dose dependency, and the *Sle1* lupus genotype, supporting a role for autoantibodies in its induction.

Based on the observed high frequency of pregnancy complications in Sle1.huTLR8tg mice, we hypothesized that analysis of early gestational timepoints, embryonic implantation and fetal placental development, would reveal premonitory changes in decidualization and placentation in Sle1.huTLR8tg pregnant mice. We found that pregnancy complications in Sle.huTLR8tg mice occurred after implantation. Although morphologic and gene expression abnormalities were present in all placentas of affected pregnancies, resorptions, placental infarctions, and fetal death occurred in a stochastic fashion and independently of fetal sex.

Toll-like receptors (TLRs) are innate immune sensors that recognize pathogens via conserved molecular patterns. Several TLRs, including TLR3 and TLR9, have been implicated in adverse pregnancy outcomes via their recognition of mitochondrial nucleic acids derived from infectious organisms or cellular damage (48, 49). TLR7 (both mouse and human) and human (hu)TLR8 are endosomal ssRNA sensors whose overexpression induces fatal SLE in mouse models. Although both TLR7 and huTLR8 recognize ssRNA, they differ in their cellular expression and function. In mice, SLE associated with excess TLR7 is primarily B cell-driven, with a minor contribution from myeloid cells (13, 16, 50, 51). Conversely, huTLR8 is highly expressed in macrophages, myeloid DCs, and neutrophils (11), with greater function than TLR7 in neutrophils (7, 52–55). Human TLR7 agonists preferentially activate DCs and pDCs, while huTLR8 agonists selectively activate monocytes (56). Finally, human TLR7 and 8 distinguish different RNA ligands and elicit distinct cytokine profiles (7). This specificity may relate to the activity of endosomal RNase T2 and RNase2, which cleave RNA at purine-uridine residues, to produce huTLR8-specific ligands not recognized by TLR7 (57, 58). Thus, these RNA-sensing endosomal TLRs have distinct functions, even within shared cellular contexts.

Placentas of Sle1.huTLR8tg mice were characterized by thinning of the junctional zone, a region situated between the decidua and labyrinth, whose loss is known to cause intrauterine growth retardation in mouse models (59). The junctional zone contributes to proper development of the fetal-placental unit by providing hormones, growth factors, and cytokines (60). Among these, members of the prolactin family and pregnancy-specific glycoprotein (PSG)/CEACAM family (61, 62) were downregulated at E8.5. Junctional zone thinning in Sle1.huTLR8tg mice appeared to involve spongiotrophoblasts and is consistent with previous work showing that delivery of viral ssRNA to healthy pregnant mice at Day E16.5 induces pro-inflammatory cytokines, placental caspase-3 activation and cell loss localized to the placental junctional zone (63). Viral ssRNA induces apoptosis of trophoblasts *in vitro* via secretion of Type 1 IFNs in a TLR8/MyD88 dependent manner (63, 64), providing further evidence of a role for endosomal RNA sensors in placental injury.

Spongiotrophoblasts are a major source of Psg16, Psg21 and Psg23 (37) all of which were downregulated in Sle1.huTLR8tg placentas. PSGs bind to glycosaminoglycans and have diverse functions in pregnancy including trophoblast adhesion and migration, proangiogenic and immunomodulatory effects (65). Mouse PSGs are not detectable in maternal sera due to their high turnover rate, but human PSG levels increase during pregnancy (37). Although reduced serum levels of PSGs have been reported in some studies of human pregnancies with adverse outcomes (38–40), we found no association between PSG1 levels and outcomes in a large longitudinal cohort of SLE and APS patients. This could be due to the broad specificity of the PSG1 antibody or because there is substantial variability in PSG levels even in healthy pregnancies (66).

The placental labyrinth contains the vascular spaces that facilitate delivery of nutrients and oxygen to the developing fetus. Although the thickness of the labyrinth was not altered in Sle.huTLR8tg mice, vasculogenesis was impaired resulting in placental infarcts and tissue inflammation as a terminal event. Importantly, although TNF was highly expressed in affected placentas, TNF deficiency did not prevent fetal loss in our model, unlike what has been shown in a passive transfer model of APL antibody-induced pregnancy loss (67), perhaps because TNF deficiency from birth induces high titers of APL antibodies in the Sle1 model (68). Nevertheless, TNF is an important effector molecule induced by huTLR8. A recent clinical trial using historical controls suggests a potential benefit of TNF inhibition starting in the first trimester in human APS pregnancies (69).

Potential mechanisms for huTLR8 induced placental injury in our model are shown in **Supplementary Figure 12**. The decreased junctional zone thickness found in Sle1.huTLR8tg but not in C57BL/6.huTLR8tg.mice supports a model in which autoantibodies bound to huTLR8 ligands, such as RNA containing immune complexes or opsonized apoptotic material, are required for placental injury. Tissue remodeling in the growing placenta may also generate a large burden of huTLR8 ligands that can bind to autoantibodies. Since huTLR8 ligands can activate monocytes and induce glycolysis in huTLR8tg bone marrow derived macrophages (23) we propose that injury is initiated or amplified by infiltrating macrophages that comprise 20–30% of leukocytes at the maternal/fetal interface (70). This hypothesis is supported by our finding that placental huTLR8tg Ly6c^+^ (classical) monocytes were preferentially activated in bone marrow chimeras, exhibiting an antigen presentation signature. We have previously shown that Classical 2 monocytes derive from infiltrating Classical 1 monocytes in lupus kidneys in both mice and humans and display a damage-associated phenotype associated with tissue injury (35). These cells were present in placental inflammatory foci together with an expanded population of CD8 T cells whose functional significance needs to be addressed in future studies. The high IFNγ expression by these cells at E9.5 may drive placental overexpression of IFNγ-inducible Gbp2b and Gm4841, which further promote inflammatory macrophages (71). Although huTLR8 is expressed in neutrophils, we did not detect an increase in neutrophil NETting of the transgenic neutrophils as an amplifying factor in this model.

APL antibodies may also directly contribute to injury in our model. They can activate TNF release from human monocytes in a huTLR8 dependent manner (21) and may induce translocation of TLR7/8 into the endosome of human monocytes and plasmacytoid dendritic cells (72). TLR expression varies throughout pregnancy (49), and huTLR8 is upregulated in inflammatory conditions including in trophoblast cells. Immunohistochemical analyses of human placentas have indicated that huTLR8 is expressed in amnion epithelial cells, chorionic trophoblasts and decidual cells with weaker expression of TLR7 and TLR9 (40, 73). Placental explants can respond to the relevant agonists in vitro by releasing inflammatory cytokines (73, 74) with a more pronounced response to huTLR8 agonists than to TLR7 agonists (73). Furthermore, *in vitro* studies using human trophoblasts have suggested that APL antibodies induce miRNAs that serve as huTLR8 ligands, resulting in release of inflammatory cytokines, neutrophil recruitment, and amplification of trophoblast injury (72). Our spatial transcriptomic analysis indicated that most huTLR8-expressing cells in the decidua and junctional zone are macrophages, closely associated with NK cells. However, there were also scattered huTLR8 expressing trophoblast cells, both in the decidua and labyrinth.

Another potential mechanism contributing to fetal loss in huTLR8tg mice involves a defect in placental NK cells. Decidual NK cells are a prevalent leukocyte population during early pregnancy, contributing to both immune tolerance (75) and to vascular remodeling. In rodent models, NK cell deficiency attenuates spiral artery remodeling and compromises fetal growth (76). Uterine NK cells also play a wide range of physiologic roles, including defense against invading pathogens, and removal of senescent decidual cells (76, 77).

Loss of NK cells in Sle1.huTLR8tg fetal-placental units was evident early in pregnancy and affected both circulating and tissue derived NK cells, suggesting a growth factor insufficiency. Among known growth factors for uNK cells, IL15 was significantly decreased in Sle1.huTLR8tg fetal-placental units by Day E8.5, corresponding to the peak time of IL15 production. IL-15, produced both by macrophages and decidual stromal cells, is essential for healthy pregnancy both in mice and humans. In our model, a decrease in IL15 mRNA expression by both cell types suggests an extrinsic defect. The regulation of IL15 transcription in placentas is not well defined and may be influenced by changes in the hormonal microenvironment (78). We did not observe any differences in the expression of Hand2 or Gata2, transcription factors known to regulate placental IL15 expression (78), in the Sle1.huTLR8tg fetal-placental units but cell type specific expression analyses will need to be performed in future experiments.

Our study has several limitations. First, although human and mouse pregnancy share several key features, there is an incomplete concordance of developmental events between mouse and human placentas. Nevertheless, a scRNAseq study of human first trimester obstetric APS placentas revealed an increase in MHC^hi^ macrophages, loss of decidual NK cells, loss of vascular endothelial cells, presence of IFNγ producing T cells, and high IFNγ, IFNβ and TNF responses, similar to what we report here (79). Second, an incomplete understanding of IL15 transcriptional and translational regulation in the placenta limits our ability to definitively determine the cause of IL15 deficiency in our model; this will be addressed in future studies. Finally, the precise mechanistic link between myeloid cell activation, CD8 T cell recruitment in the maternal layers, and the subsequent loss of the junctional zone remains to be elucidated.

Nevertheless, this model provides a valuable platform to investigate early events contributing to APL antibody-induced pregnancy loss, which are challenging to address in a clinical setting. A deeper understanding of these early mechanisms will facilitate the identification of at-risk pregnancies and the development of targeted therapeutic strategies to protect both mother and fetus.

## Materials and Methods

### Mice

Clone 8 huTLR8 bacterial artificial chromosome (BAC) transgenic mice (11) on the C57BL/6 background were a kind gift of C. Guiducci (Dynavax). Clone 8 mice were bred to Sle1.Yaa mice and were maintained either as homozygous (*Ho*) or heterozygous (*Ht*) Sle1.huTLR8tg. SLE1 genotyping for colony maintenance was performed using primers shown in **Supplementary Table 8**. Mice were typed for hetero– or homozygosity for the huTLR8 transgene as previously described (23). Genotyping of fetal tissue for the Y chromosome was performed by Transnetyx. For harvest of organs, mice were perfused with 60cc of PBS. To distinguish circulating cells from resident immune cells in tissue mice were injected retro-orbitally with 0.5ug of PE conjugated antibodies to CD45.2 or CD45 (eBIOscience) depending on the analysis panel 10 minutes prior to perfusion. Only female mice were used.

### Quantitation of Autoantibodies to Cardiolipin, Sm/RNP and Chromatin

ELISA assays for antibodies to CL/β2GP1, Sm/RNP, chromatin and dsDNA were performed on serial serum samples as previously described (80–83) using sera dilutions of 1/250 for anti-CL/β2GP1, –Sm/RNP and –dsDNA and 1/500 for anti-chromatin. A high titer serum was run in serial dilutions on each plate. Using this control as a standard, OD values were converted to arbitrary units using regression analysis (Prism Version 9). Maximal units set for each ELISA correspond with the test serum dilution and represent the OD at which the plateau for the ELISA is reached.

### Analysis of Pregnancy Outcomes

Pregnancies were monitored and pregnant females displaying displayed distress due to dystocia were euthanized. Pups were weighed and genotyped for gender, and placentas were harvested for further analysis. For timed analyses, the day of vaginal plug detection was defined as E0.5, and uteri were harvested at E8.5-9.5, E13.5 or E18.5.

### Placental Histology/ Immunopathology/ Immunohistochemistry

Placental tissue was fixed in 10% formalin for 24 hours, then transferred to 70% ethanol solution followed by embedding in paraffin. All embedding, sectioning, and immunohistochemical staining were performed by Histowiz, NY, and digitized images were provided for analysis. Tissue was sectioned through the center and stained with H&E, and with antibodies to MPO to detect neutrophils and SMA to stain blood vessel walls. Dolichos biflorus agglutinin (DBA) lectin (Vector Labs, 1:1000 dilution) was used to detect uterine natural killer cells. Periodic acid–Schiff (PAS) stain was used to detect glycogen-containing cells in the placental junctional zone. Thickness of the placental layers on PAS-stained images was measured using a digital caliper and reported as the average of 3 measurements across the placenta. NK cell density was assessed by averaging the number of NK cells in 3 contiguous areas of the decidua each measuring 0.25μm^2^. Placental arteries were identified (4-5 per placenta), and luminal and wall diameters were measured using ImageJ software. SMA score was measured on a semi-quantitative scale of 0-3 by a reader blinded to the maternal genotype.

### Placental RNA Isolation and RT-qPCR

Placental RNA was isolated using RNeasy isolation kit (Qiagen) according to manufacturers’ protocol with on column DNAse I digestion. RNA quality and concentration were assessed using Bioanalyzer. cDNA was synthesized using the Superscript III First-Strand Synthesis kit with random hexamer protocol according to manufacturers’ protocol, including an RNAse H step (Thermo Fisher Scientific). RT-qPCR was performed using Roche Lightcycler® 480 SYBR green master I and primers for βactin, TNF and IL-6 as previously described and for MPO and developmental genes as shown in **Supplementary Table 8.** Data was analyzed using the ΔΔCT method.

### Multiparameter Flow cytometry

Pregnant mice were perfused with PBS to eliminate circulating leukocytes. Minced kidneys and placentas were incubated in 500ug/mL Liberase (Liberase TL, Roche) and 100U/mL DNase I (Roche) for 15 min at 37°C. Tissues were then filtered through a 70μM filter, washed with DMEM (Gibco) and resuspended in FACS staining buffer (2.5% FBS, 0.05% sodium azide in PBS). Heparinized blood was sampled via cheek bleeding pre– and post-infusion and lysed using 1x BD Pharm Lyse buffer (BD Pharmingen). Spleens and bone marrows were mechanically disrupted. Cells were stained in FACS staining buffer followed by Fc-block for 10 min at 4C and fluorochrome-conjugated antibody panels as previously described (68, 84) (**Supplementary Table 9**). Fixable viability dye (eFluor780; eBioscience™ Thermo Fisher Scientific) was used to exclude dead cells from analysis. All samples were acquired on BD Symphony or BD Fortessa flow cytometer (BD Biosciences) and analyzed with FlowJo software (Tree Star).

### Assessment of NETS

Bone marrow neutrophils were isolated from 3– and 6-month-old homozygous female Sle1.huTLR8tg, Sle1 and C57BL/6 controls using EasySep mouse neutrophil enrichment kit (STEMCELL) and were treated with PMA 50nM for 4 hours at 37°C with or without pre-treatment with PMA or TLR8 agonist CL075 (Invivogen) 10mg/ml for 20 minutes. NETs were detected using flow cytometry after staining for MPO and citrullinated histone H3 (**Supplementary Table 9**).

### RNA Sequencing

RNA from whole fetal-placental units (E8.5-9.5) or whole placentas (E13.5), was isolated using Qiagen mRNeasy kit and subjected to bulk RNA sequencing (Novogene). Sequences were aligned to the mm10 genome with STAR aligner (85) and the read count for each transcript was extracted from the alignment with HTSeq program (86). Transcripts with <100 reads across all samples were filtered out. The read counts were normalized to the same total read counts level at sample level, log2 transformed, and then quantile normalized.

Differential gene expression analyses were carried out by LIMMA test (87). Differentially Expressed Genes (DEGs) were identified at p-value <0.05. Biological functional/pathways enriched for DEGs were determined by fisher-exact test at p value <0.05 using the biological process categories in Gene Ontology (GO) (88) and pathways curated in several pathway databases (KEGG, Ingenuity IPA, BIOCARTA, NABA, Panther, PID, REACTOME, Wiki-pathway). Normalized RNAseq data were also used to run Gene Set Enrichment Analysis (GSEA) using the Broad Institutes’ GSEA 4.0.3 software (89). Genes were run against three gene sets from Gene Ontology (GO): GO:0007565∼female pregnancy, GO:0019835∼cytolysis, and GO:0048002∼antigen processing and presentation of peptide antigen. Data is available on Geo (GSE316139).

### Bone Marrow Chimeras

8– to 10-week-old Sle1.CD45.1 mice were treated with biotin anti-CD117 and streptavidin-saporin as previously described (36) and 2 x 10^6^ CD45.2 bone marrow cells from Sle1 or Sle1.huTLR8tg donors were transferred i.v. 8 days later. Mice were monitored for chimerism every 2 weeks using flow cytometry of peripheral blood and were mated 6-8 weeks after bone marrow transfer.

### PSG1 ELISA

Demographic and outcome data from the PROMISSE study have been previously published (18). ELISA was performed using a commercial kit (Quantikine – R&D Systems) according to manufacturers’ instructions using the provided quantitation control. Sera were diluted from 1:250 to 1:10,000 depending on the pregnancy stage. Data are expressed as ng/ml.

### Single cell RNA sequencing

To obtain single cell suspensions, 3-5 pooled placentas from 5 individual E9.5 Sle1 and 5 Sle1.huTLR8tg uteri were incubated for 15 minutes in DMEM containing 1 mg/ml Collagenase D (Roche), 100U/ml DNAse I (Sigma), 0.25mg/ml Liberase (Invitrogen) at 37°C. Tissues were gently dissociated using a pipette and then passed through a 70 μm cell strainer (BD). The single cell suspension was stained with Live-Dead stain and anti-CD45 and single CD45+ and CD45-fractions were flow sorted on a BD FACS Aria II and mixed at a 30:70 ratio then immediately fixed using the 10X Genomics GEM-FLEX V2 preparation kit and frozen until use. After all samples had been collected, cells were thawed and washed. 5-10 x 10^3^ cells per sample were loaded onto the 10X Genomics GEM-X mouse 4-plex chip and processed per manufacturer’s instructions. DNA amplification and library construction were carried out as per manufacturer’s instructions and pooled sequencing was performed using Novaseq S1.

Alignment and quantification were performed using Cell Ranger v7.1.0. Analysis of the resulting data was done using Seurat v4 (90). Batch correction was performed using Harmony (91). For clustering of CD45+ cells, we took a step-wise approach, starting with coarse clustering to identify the main immune lineages (T cells, NK cells, B cells, myeloid cells). We focus here on the myeloid cell lineage which was then clustered separately. Analysis of other data will be reported separately. Cluster labeling was done by expression of canonical markers and by reference mapping to our previously published renal scRNA-seq data (35), while the clustering resolution was chosen to obtain clusters that are relatively uniform in their predicted reference identities (**Supplementary Table 10**).

### Spatial Transcriptomics

Murine placentas from 3 E9.5 Sle1 and 5 Sle1.huTLR8tg mice were processed for Xenium spatial transcriptomics according to the manufacturer’s protocol and characterized using both custom and manufacturer-provided multi-tissue probe panels, as previously described (35) with the addition of probes for IL15. Spatially resolved gene expression data were processed and analyzed using Seurat and Xenium Explorer (version 3.1). Cells in the lowest 10% for UMI and gene counts were excluded. Data normalization and feature selection were performed using the NormalizeData and FindVariableFeatures functions in Seurat. We used a standard Seurat pipeline (filtering low quality cells, normalization, scaling) to identify 26 clusters by K-means clustering representing tissue and immune cells; full cluster annotations will be reported elsewhere. Placental stromal clusters in the mLAP and decidua were identified based on coexpression of Wt1 and Pdgfra. Myeloid cells were defined by (i) manual curation to remove non-myeloid cells, proliferating cells, and doublets, followed by (ii) assignment of myeloid identities using the scRNAseq reference described above; here, we integrated the spatial and single cell myeloid data sets with the FindIntegrationAnchors and FindTransferAnchors functions in Seurat. This integration enabled annotation of cells in the Xenium (query) dataset based on the reference while preserving spatial coordinates. To localize Xenium myeloid subsets with high correspondence to the single-cell reference, we mapped cells with prediction scores > 0.75 in situ using Xenium Explorer (version 3.0). We quantified the number of transcripts of huTLR8, IL15, CD4, CD8, and IFNG per cell type normalized per mm² of tissue in each sample using Xenium software and Prism. To map inflammatory aggregates, we calculated the fraction of cells in each cluster relative to total myeloid cells within six inflammatory aggregates from five Sle1.huTLR8tg placentas of the kidney.

## Statistical Analyses

Pairwise comparisons were analyzed using non-parametric Mann Whitney-U test to compare medians, and normally distributed data were analyzed by an independent t-test to compare means. Multiple group comparisons were analyzed using non-parametric Kruskal Wallis ANOVA with Dunn’s correction for multiple comparisons. For correlation studies, the Spearman’s rho (rs) or Pearson correlation (rp) were calculated for non-parametric and normally distributed data, respectively. Proteinuria and survival data were analyzed using Kaplan-Meier curves and the log rank test. PSG1 association with pregnancy outcomes was evaluated using a mixed model. Values of p<0.05* were considered statistically significant. Only significant *P* values are shown. Statistical analysis was performed using GraphPad Prism.

## Study approval

All experiments were performed under protocols approved by the Institutional Animal Care and Use Committee (IACUC) of the Feinstein Institutes for Medical Research

## Data availability

RNAseq data is available on Geo (GSE316139). Supporting data are included in a Supplementary file.

## Author Contributions

NM: designed research studies, conducted experiments, acquired data, analyzed data, co-wrote the manuscript

YX: designed research studies, conducted experiments, acquired data, analyzed data, co-wrote the manuscript

### Co-first authorship in alphabetical order

CR: conducted experiments, acquired data, analyzed data

KL: conducted experiments, acquired data, analyzed data

SM: conducted experiments, acquired data

ZY: analyzed data

WZ: analyzed data

MA: designed research studies, conducted experiments, acquired data

PW: designed research studies

MG: designed research studies, acquired data

JSa: designed research studies, acquired data

JSo: designed research studies

AA: acquired data, analyzed data

PH: acquired data, analyzed data

AD: designed research studies, acquired data, analyzed data, co-wrote the manuscript

## Acknowledgements

We thank Adnan Arif, MD and Haiou Tao, BSc for technical assistance.

