## Supplementary Figures for "Pregnancy loss due to early developmental defects in lupus mice expressing human TLR8"

**Supplementary Figure 1. Splenic B cell subsets in female B6.huTLR8tg (grey), Sle1 (blue) and Sle1.huTLR8tg (red) mice at sequential ages.** **A, B.** CD19 B cell number (A) and percent (B). **C, D.** Follicular B cell number (C) and percent of CD19 B cells (D); **E, F.** Marginal zone B cell number (E) and percent of CD19 B cells (F); **G, H.** Germinal center B cell number (G) and percent of CD19 B cells (H); **I, J**. Plasma cell number (I) and percent of spleen cells (J); Each symbol represents an individual spleen; lines represent the median. p values calculated for age-related changes in each strain and strain specific differences using Kruskal Wallis ANOVA with Dunn's correction for multiple analyses. *p<0.05, **p<0.01, ***p<0.001.

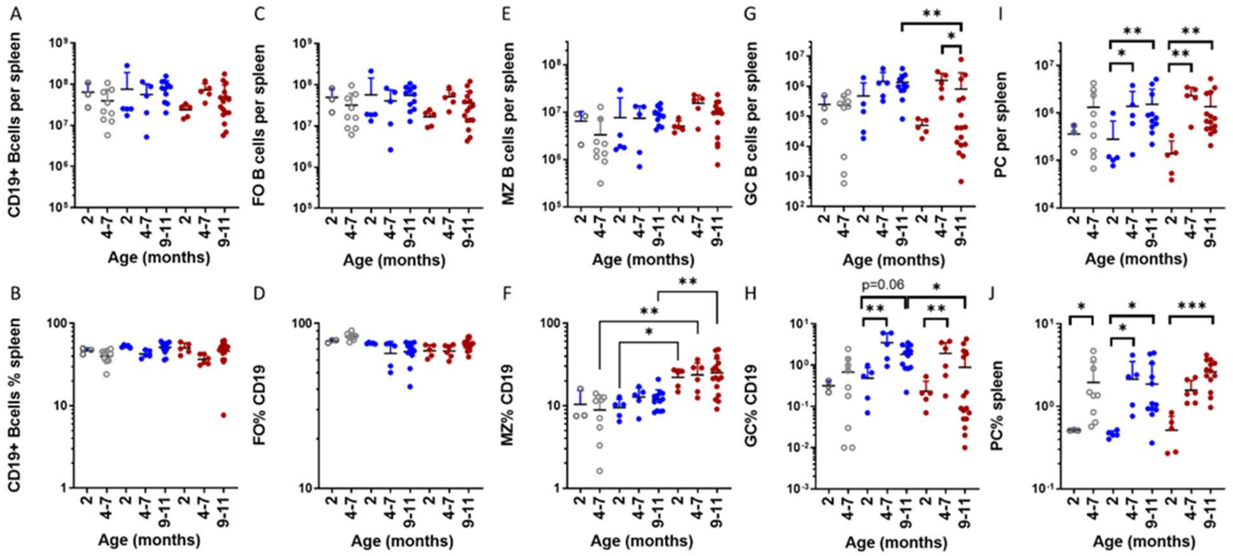

**Supplementary Figure 2. Splenic myeloid and T cell subsets in female B6.huTLR8tg (grey), Sle1 (blue) and Sle1.huTLR8tg (red) mice at sequential ages. A.** Total cell count per spleen; **B, C.** CD11b+ percent (B) and number (C): **D, E.** CD4 T cell percent (D) and number (E); **F, G.** CD8 T cell percent (F) and number (G); **H, I.** TFH as a percent of CD4 T cells (H) and total number per spleen (I); Each symbol represents an individual spleen; lines represent the median. p values calculated for age-related changes in each strain using Kruskal Wallis ANOVA with Dunn's correction for multiple analyses. *p<0.05, **p<0.01. **J.** Representative immunohistochemistry of spleens from 9-month-old Sle1 (upper) and Sle1.huTLR8tg (lower) mice showing loss of GL7+ germinal centers in Sle1.huTLR8tg mice.

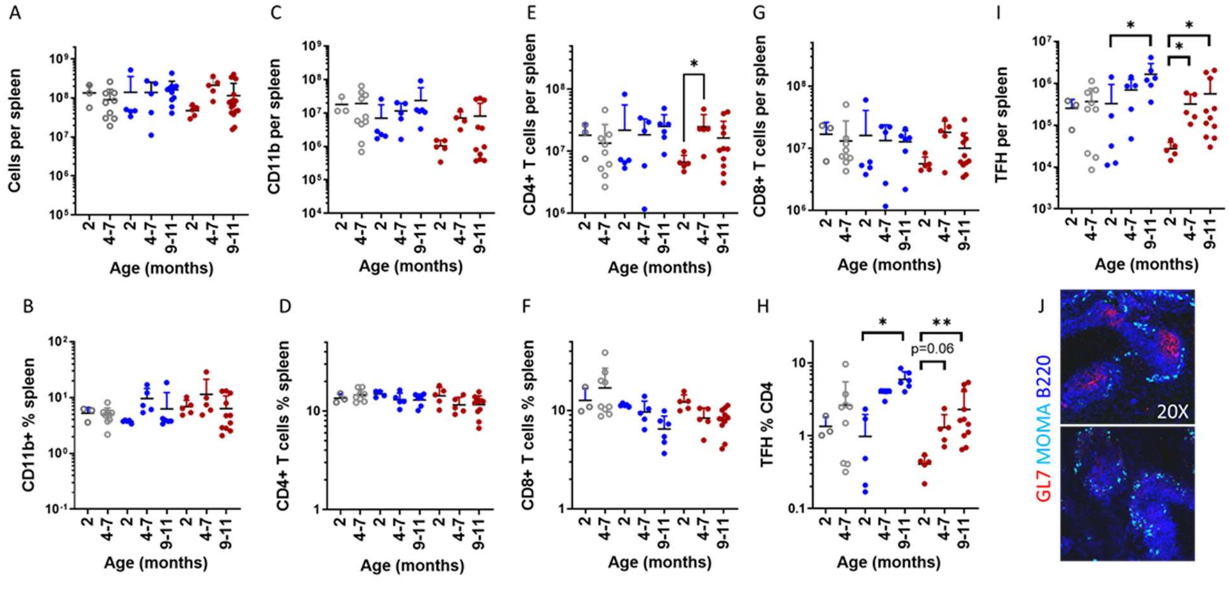

**Supplementary Figure 3. Placental inflammatory markers. A, B**. qPCR of myeloperoxidase (MPO; A) and TNF (B) expression in placentas from mice of the indicated genotypes. Each symbol represents an individual placenta; lines represent the median. p values calculated using Kruskal Wallis ANOVA with Dunn's correction for multiple analyses. *p<0.05, **p<0.01, ****p<0.0001. **C, D**. Negative correlation of MPO (C) and TNF (D) expression with placental weight. p values calculated using simple linear regression.

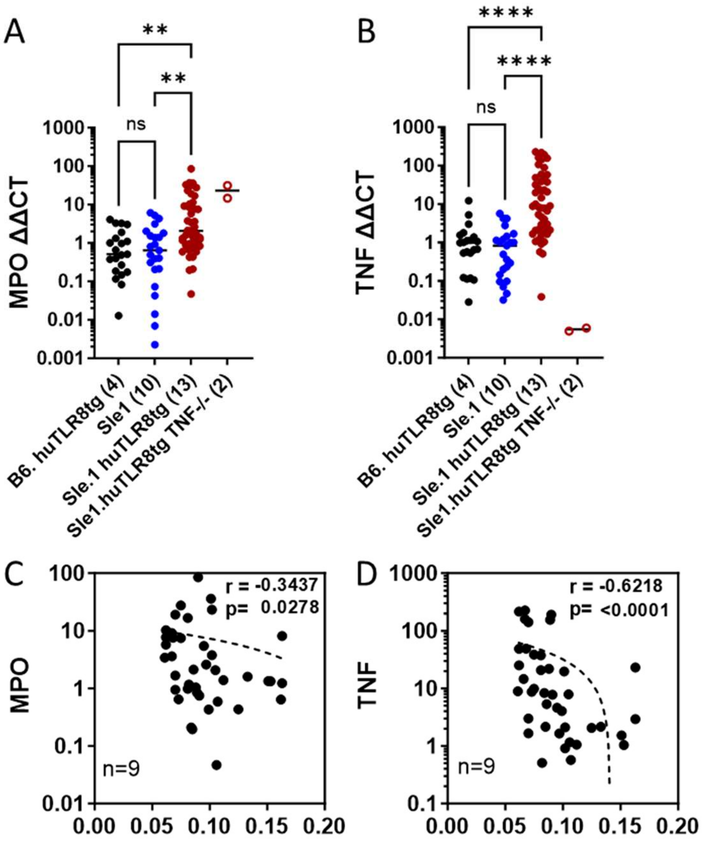

**Supplementary Figure 4**. **Evaluation of neutrophil NETting**. **A**. Gating strategy for bone marrow-derived neutrophils stimulated to produce NETs. After gating on size, singlets and Ly6G, cells stimulated with the indicated conditions were evaluated for staining for myeloperoxidase (MPO) and citrullinated histone 3 (Cit H3). **B**. Percent NETting neutrophils in female C57BL/6, Sle1 and Sle1huTLR8tg mice at 2-3 months (black circles) and 6 months (grey circles) of age after stimulation with the indicated conditions. Each symbol represents an individual mouse. Comparisons between strains showed no differences.

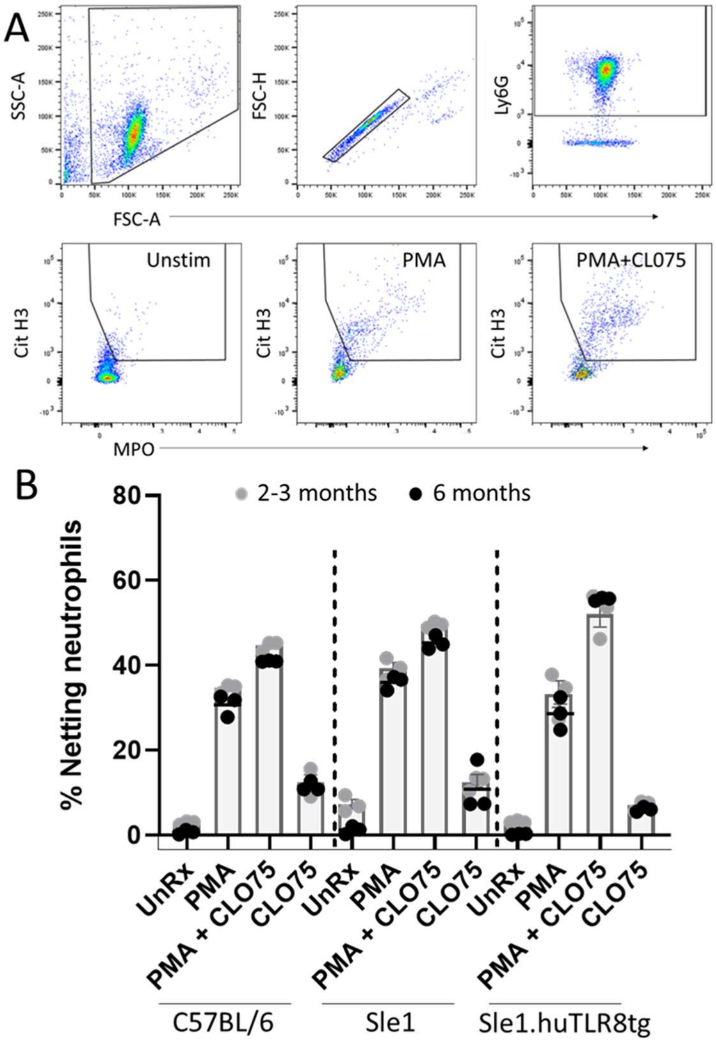

**Supplementary Figure 5. Evaluation of markers of preeclampsia. A, B.** qPCR of selected genes from Day E8.5 (A) fetal-placental units and Day E13.5 (B) placentas. **C.** Systolic and diastolic blood pressure in two Sle1 and three Sle1.huTLR8tg mice measured longitudinally throughout gestation. **D.** sFlt1 serum levels at term. **E.** Serum VEGF levels from non-pregnant mice **(**E0) and at E9.5 show no difference between Sle1 and Sle1.huTLR8tg mice. A, B, D, E. Statistics calculated using non-parametric Mann Whitney t-tests for each analyte. *p<0.05.

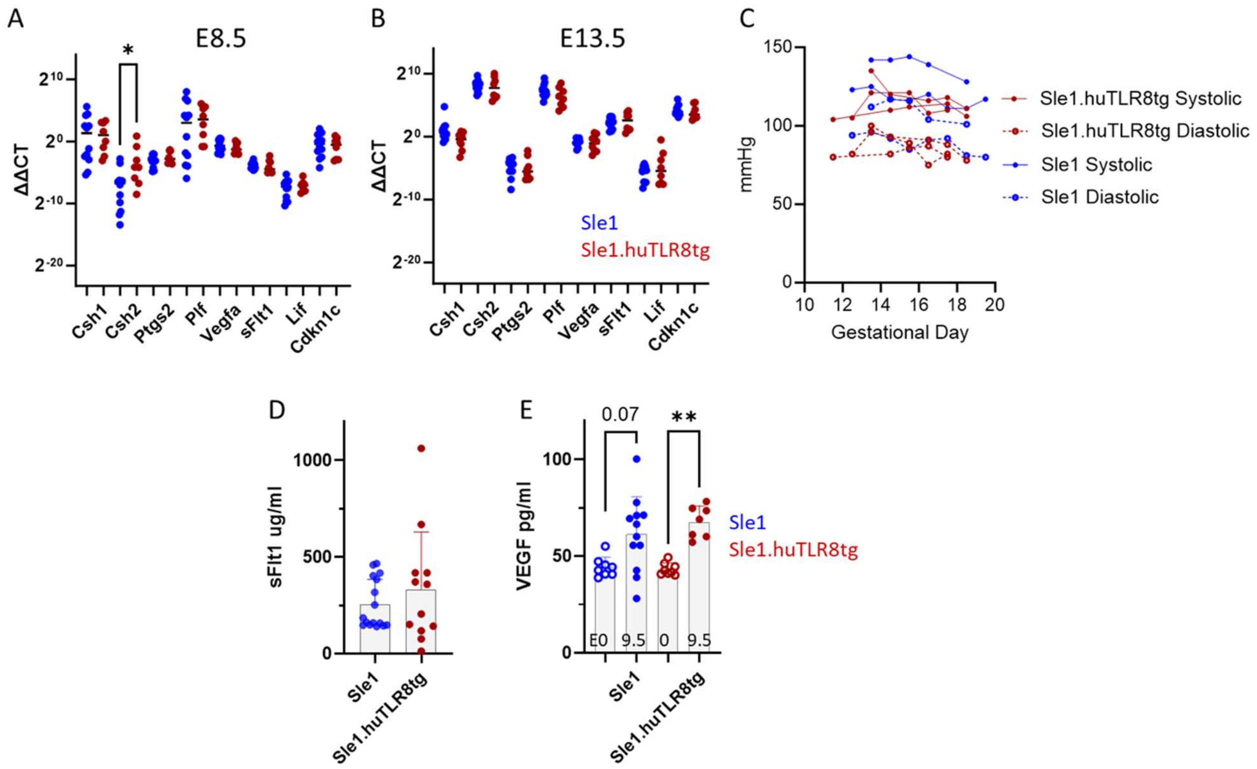

**Supplementary Figure 6. Analysis of junctional zone cells**. **A, B**. No difference in the thickness of the primary trophoblast giant cell layer between strains at E9.5. **C**. Representative images from E13.5 Sle1 (i, iii) and Sle1.huTLR8tg (ii, iv) placentas show junctional zone thinning and decreased DBA positive cells in Sle1.huTLR8tg placentas (enumerated in Figure 5D). Trophoblast giant cells (blue arrows) and glycogen cells (white arrows) do not differ between strains. A, C: Placentas stained with PAS and DBA. Representative of 3-5 placentas per group.

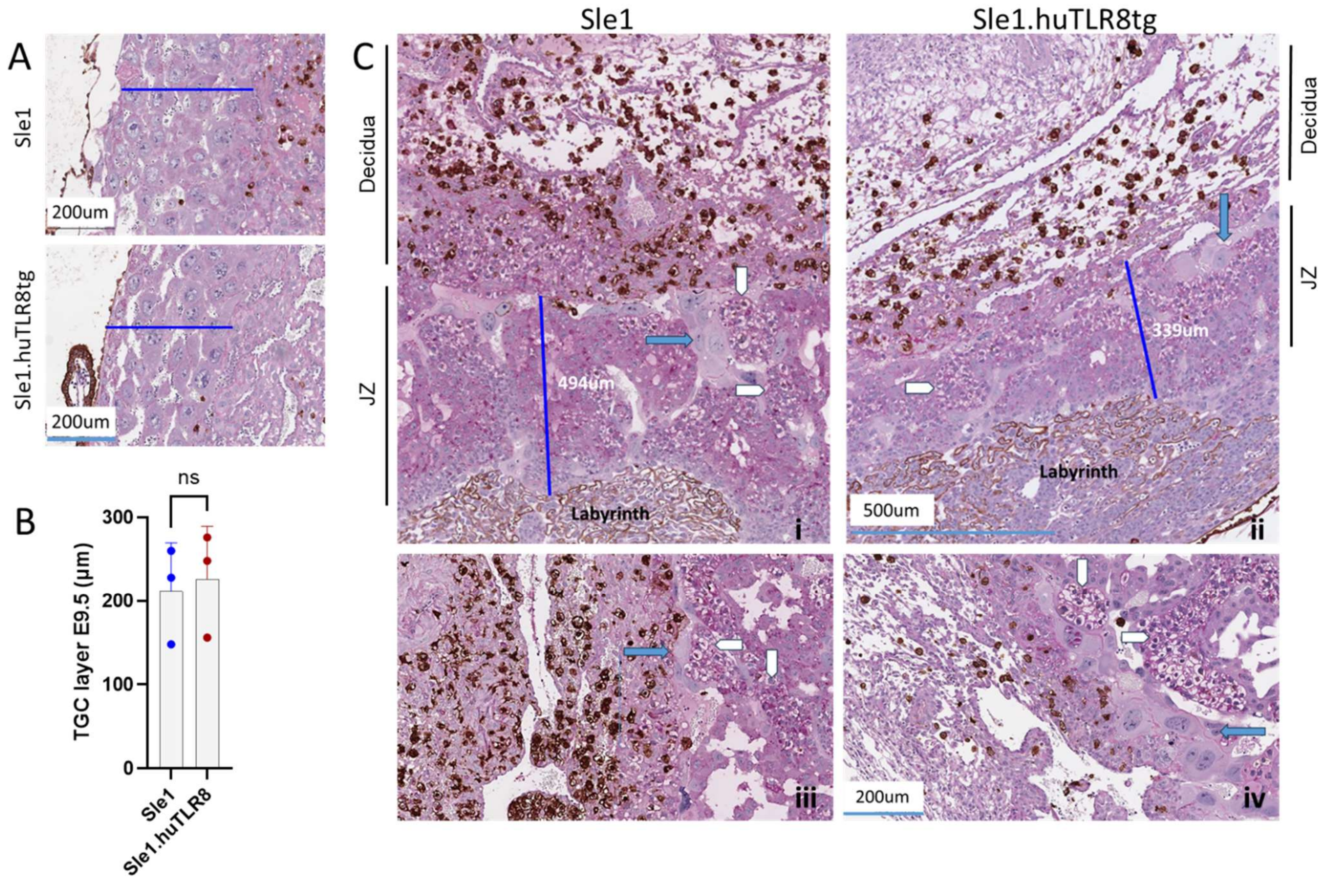

**Supplementary Figure 7**. **Gating strategy for placental NK cells and CD8 T cells**. **A.** Live singlets were gated for $CD45^{+}/CD45.2^{-}$ cells to exclude intravascular cells (R1 gate). NK cells were defined as $NK1.1^{+}CD3^{-}CD8^{-}CD19^{-}CD11b^{-}$. Tissue resident NK cells were defined as $CD49a^{+}CD49b^{-}$ and circulating NK cells were defined as $CD49a^{-}CD49b^{+}$. **B.** Representative plots from E9.5 and E13.5 placentas show that tissue ($CD45.2^{-}$) NK cells are predominantly tissue resident NK cells and intravascular ($CD45.2^{+}$) NK cells are predominantly circulating NK cells. **C, D.** The proportion of tissue resident (tr) and circulating (c) uterine NK cells does not differ between strains at Day E9.5 (C) or Day E13.5 (D). **E, F.** The frequency of CD8 T cells is increased in Sle1.huTLR8tg placentas at E9.5 (E) with a continuing trend at E13.5 (F). C-F: Each symbol represents a placental pool from a unique uterus. Bars represent the mean + SD. p values calculated using Kruskal Wallis ANOVA with Dunn's correction for multiple analyses. $*p<0.05$.

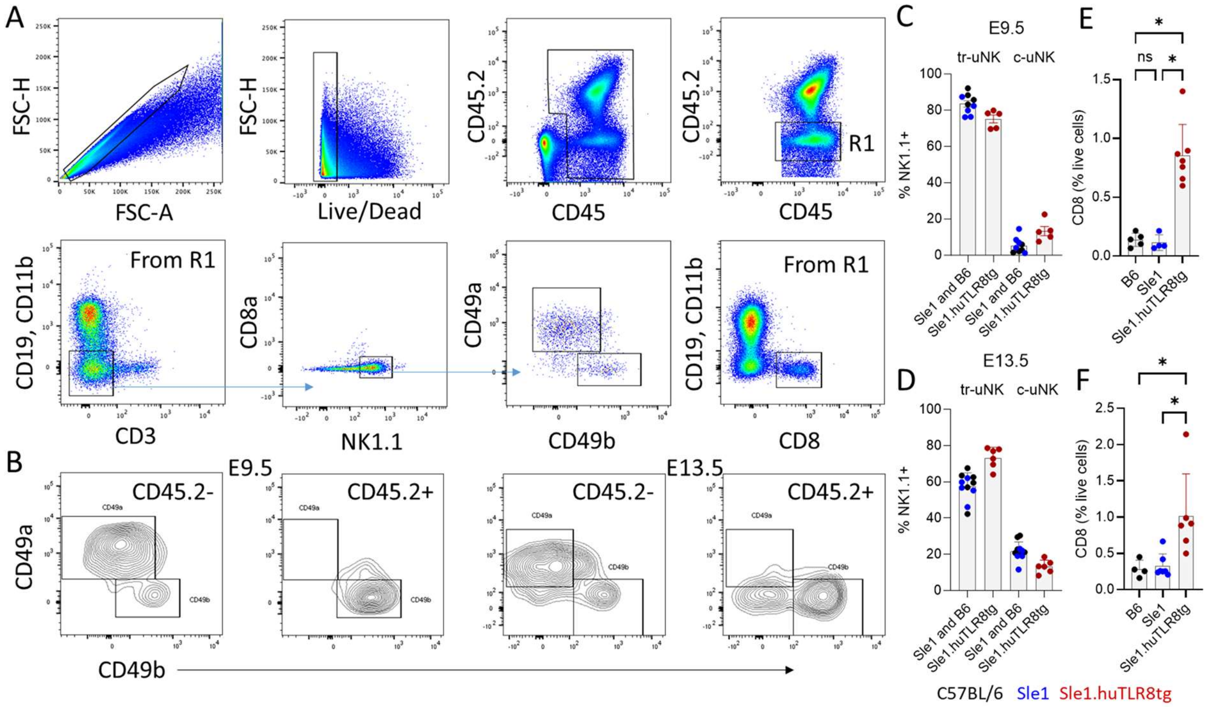

**Supplementary Figure 8**. **Gating strategy for placental myeloid cells. A.** Live $CD45^+/CD11b^+$ singlets were gated for $CD45.2^-$ cells to separate tissue (R1 gate) from intravascular cells (R2 gate) and the R1 gate was further gated to separate $Ly6G^+$ neutrophils, $XCR1^+$ cDC1, $Ly6C^+$ classical monocytes (CΦ), $CD11a^+/Ly6C^-$ non-classical monocytes (NCΦ) and $CD11c^+/CD26^+$ cDC2. The remaining cells (R5) were then gated on F4/80 and CD81 to define resident macrophages with only a small proportion of cells remaining unclassified. **B.** Percent live tissue $CD11b^+$ cells is increased in E9.5 Sle1.huTLR8tg placentas. **C.** No difference in the relative frequency of CD11b+ subsets between strains. Each symbol represents a placental pool from a unique uterus. Bars represent mean + SD. p values calculated using Kruskal Wallis ANOVA with Dunn's correction for multiple analyses. D. Frequency of neutrophils is increased at term in Sle1.huTLR8tg placentas. B, D. Each symbol represents a placental pool from a unique uterus. Bars represent mean + SD. p values calculated for comparisons between each gestational age using non-parametric Mann Whitney t-tests. *p $<0.05$, **p$<0.01$.

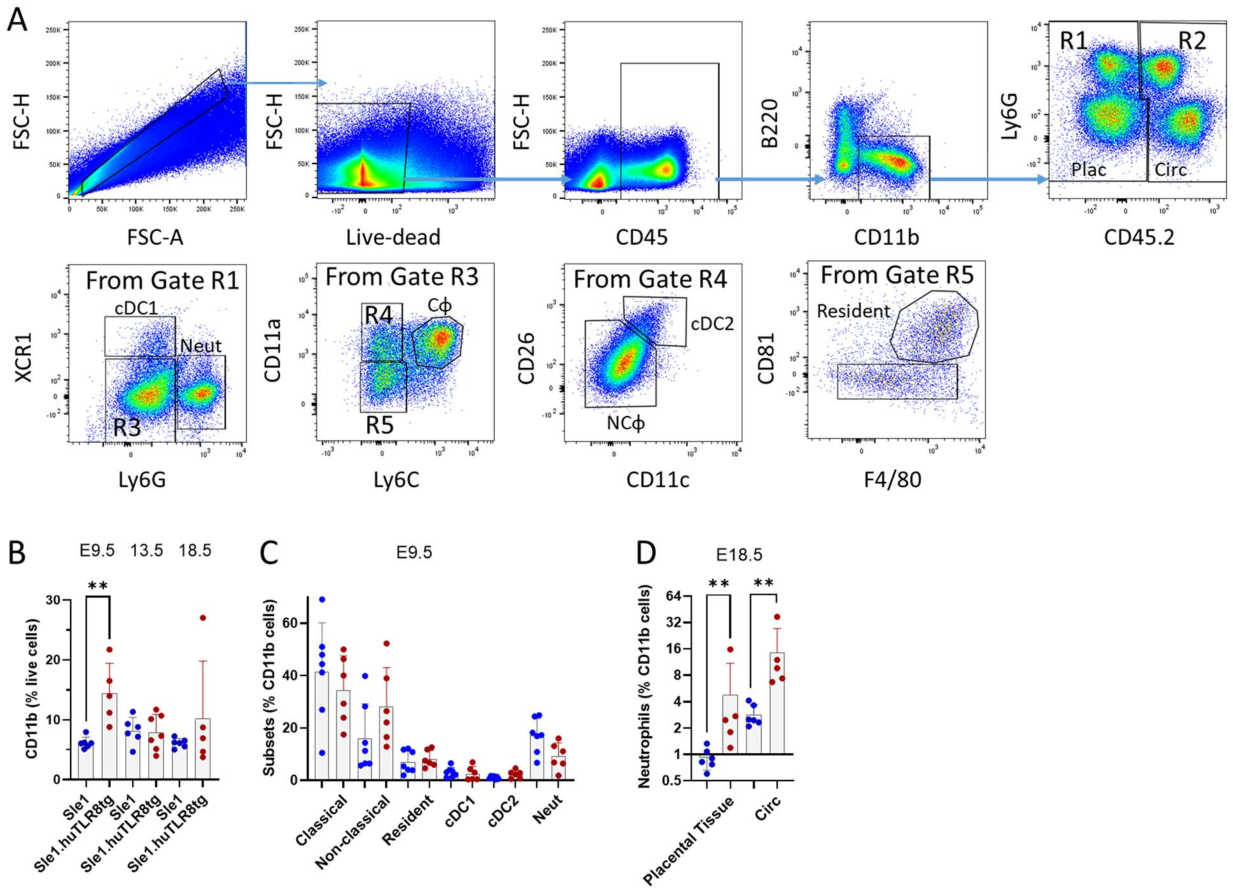
A
FSC-H
FSC-A
Live-dead
CD45
B220
CD11b
Ly6G
CD45.2
R1
R2
Plac
Circ
From Gate R1
XCR1
Ly6G
cDC1
Neut
R3
From Gate R3
CD11a
Ly6C
R4
Cϕ
R5
From Gate R4
CD26
CD11c
cDC2
NCϕ
From Gate R5
CD81
F4/80
Resident
B
E9.5
13.5
18.5
CD11b (% live cells)
Sle1
Sle1.huTLR8tg
C
E9.5
Subsets (% CD11b cells)
Classical
Non-classical
Resident
cDC1
cDC2
Neut
D
E18.5
Neutrophils (% CD11b cells)
Placental Tissue
Circ

**Supplementary Figure 9**. **Analysis of myeloid cell enrichment in bone marrow chimeras. A.** Live $CD45^-/CD11b^+$ (tissue) and $CD45^+/CD11b^+$ (intravascular) cells from placentas or total $CD11b^+$ cells from blood were gated on CD45.1 and CD45.2 to separate donor (CD45.2) from recipient (CD45.1) cells. **B.** The percentage of donor cells in the placenta was compared with that in the blood to detect enrichment of donor cells in the placenta (a ratio of 1 indicates no enrichment). Myeloid subsets were classified as in Supplementary Figure 8. Bars represent mean + SD. **C.** $Ly6C^{hi}MHCII^{hi}$ myeloid cells are increased in the placentas compared with blood with no difference between strains. Each symbol represents a placental pool from a unique uterus. Lines indicate the median. p values calculated using Kruskal Wallis ANOVA with Dunn's correction for multiple analyses. *$p<0.05$. **D.** Enrichment of Sle1.huTLR8tg donor cells among the $Ly6C^{hi}$ $MHCII^{hi}$ myeloid subset in both blood and placenta. Each symbol represents a placental pool from a unique pregnancy. Bars represent mean + SD. p values for each tissue calculated using Mann-Whitney t-test. *$p<0.05$. **E.** Gating strategy for D after identifying $Ly6C^{hi}$ myeloid cells as in Supplementary Figure 8 – a representative Sle1 mouse is shown. **F, G.** Increased MFI of CD86 in the placental $Ly6C^{hi}$ $MHCII^{hi}$ myeloid subset compared with the $Ly6C^{hi}$ $MHCII^{lo}$ subset in Sle1.huTLR8tg but not in Sle1 placentas. Representative histograms are shown in (G). Each symbol represents a placental pool from a unique uterus. Bars represent mean + SD. p values for each strain calculated using paired t-test. **$p<0.01$.

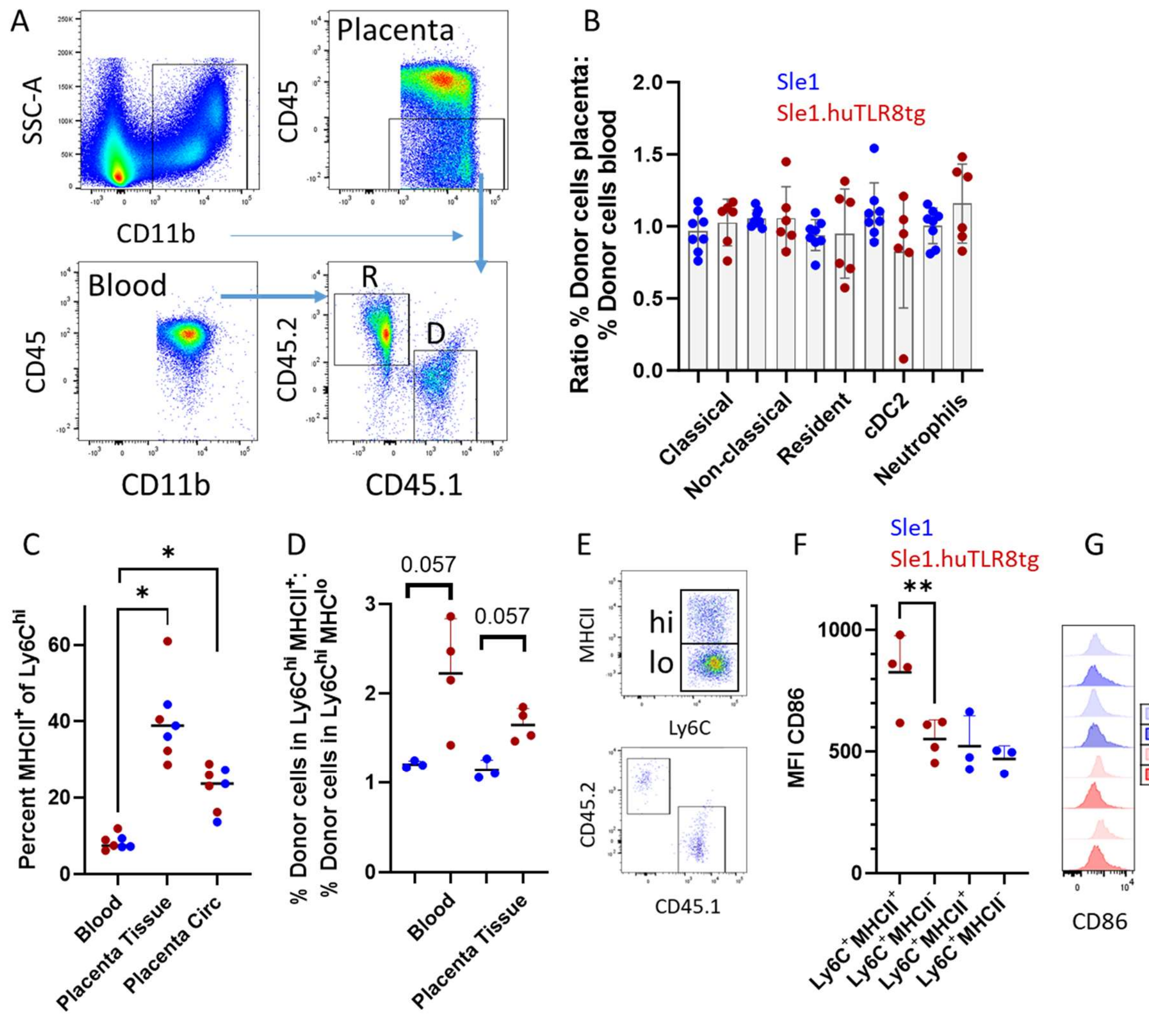

A
SSC-A
CD11b
Placenta
CD45
Blood
CD45
CD11b
R
D
CD45.2
CD45.1
B
Sle1
Sle1.huTLR8tg
Ratio % Donor cells placenta: % Donor cells blood
Classical
Non-classical
Resident
cDC2
Neutrophils
C
Percent MHCII+ of Ly6Chi
Blood
Placenta Tissue
Placenta Circ
D
% Donor cells in Ly6Chi MHCII+: % Donor cells in Ly6Chi MHClo
0.057
0.057
Blood
Placenta Tissue
E
MHCII
hi
lo
Ly6C
CD45.2
CD45.1
F
Sle1
Sle1.huTLR8tg
MFI CD86
Ly6C+MHCII+
Ly6C+MHCII-
Ly6C+MHCII+
Ly6C+MHCII-
G
MHChi
MHClo
MHChi
MHClo
CD86

**Supplementary Figure 10. Spatial transcriptomic analysis of E9.5 placentas**. **A.** Percent of myeloid and non-myeloid cells expressing ≥1 huTLR8 transcript. **B.** Representative image showing huTLR8 colocalization with Itgam. Myeloid cells are indicated in pink. **C.** Representative image showing expression of NK cell genes Ncr1 (pink dots) and Klrb1c (blue dots). **D, E.** Representative images showing IL15 transcripts (yellow dots) in myeloid cells (red outlines) or stromal cells from the mLAP (blue outlines) or decidua (orange outlines). C-E. Nuclei are stained with DAPI (white). **F-I.** Enumeration of IL15 expression shown as percent stromal (F) or myeloid (H) cells expressing ≥1 IL15 transcript and the average number of transcripts per cell (G, I). Each symbol represents an individual placenta. Bars represent mean + SD. p values for cell type calculated using Mann-Whitney t-test. *$p<0.01$. **J, K.** Representative images showing myeloid cell subsets in the uterine wall, mLAP and decidua. **L, M.** Representative images showing small clusters of myeloid cells in the decidua. J-M. Nuclei are stained with DAPI (white).

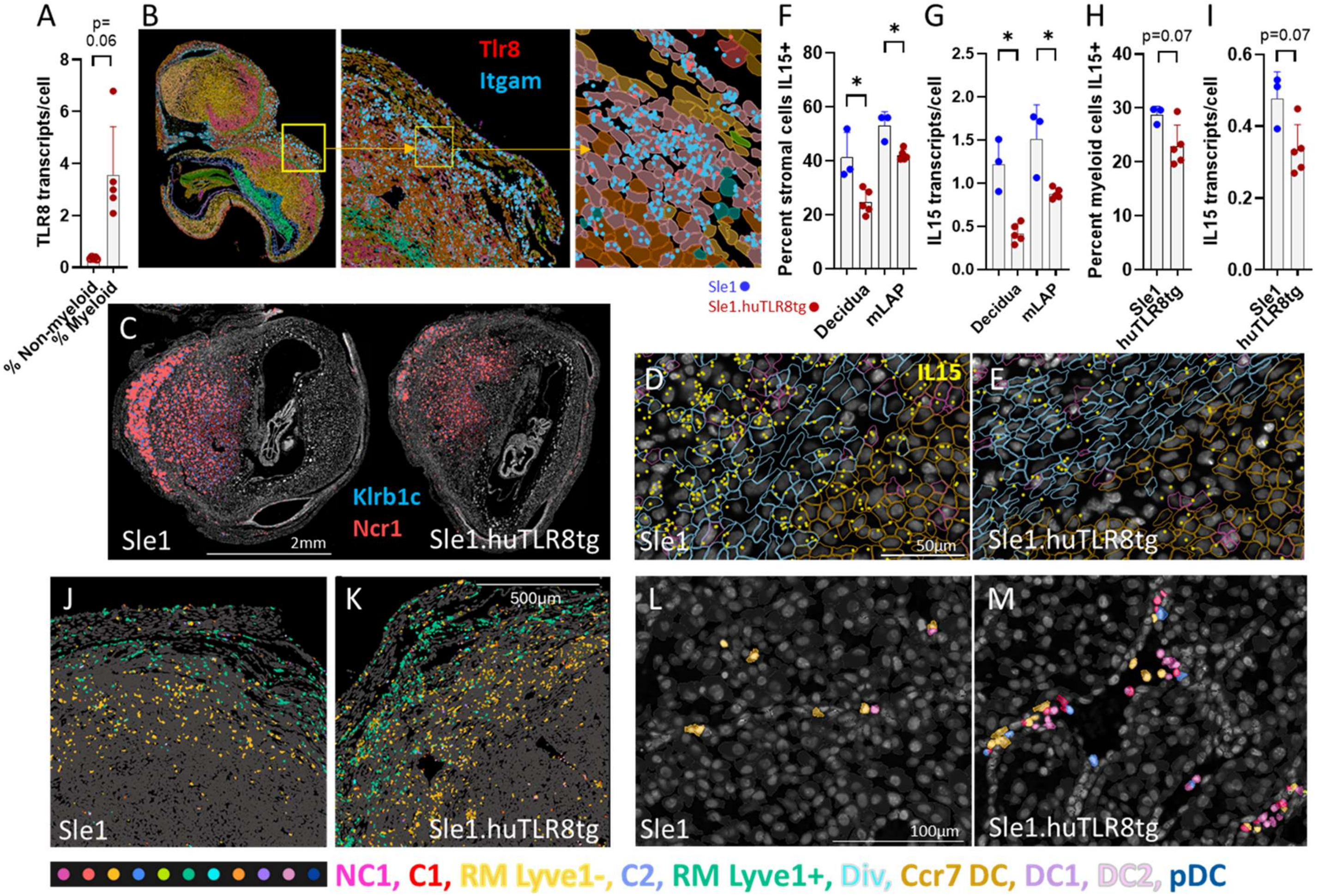

A
B
C
D
E
F
G
H
I
J
K
L
M
p=
0.06
TLR8 transcripts/cell
% Non-myeloid
% Myeloid
Tlr8
Itgam
Sle1
Sle1.huTLR8tg
Percent stromal cells IL15+
Decidua
mLAP
IL15 transcripts/cell
Percent myeloid cells IL15+
p=0.07
Klrb1c
Ncr1
2mm
IL15
50µm
500µm
100µm
NC1, C1, RM Lyve1-, C2, RM Lyve1+, Div, Ccr7 DC, DC1, DC2, pDC

**Supplementary Figure 11**. **Serum PSG1 levels. A-C.** Serum PSG1 levels in longitudinal samples from healthy donors (A) and SLE and/or APS pregnancies (B, C) in the PROMISSE cohort. Outcome in the SLE and/or APS cohorts was classified as good if there was a healthy delivery at >38 weeks or as adverse if there was IUGR, fetal demise or preeclampsia (31). No differences in serum PSG1 levels were detected across groups.

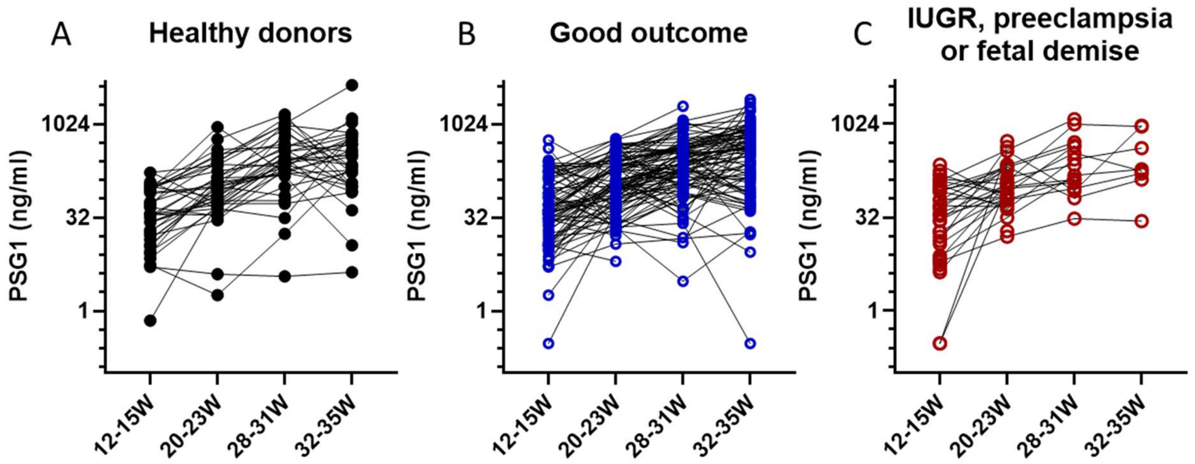

**Supplementary Figure 12**: Model of pathogenesis of huTLR8 induced pregnancy loss. A. TLR8 ligands binding to autoantibodies or autoantibodies alone may have direct cytotoxic effects on trophoblasts. B. Activation of TLR8 in myeloid cells by nucleic acid containing debris or immune complexes results in upregulation of MHC11, CD86 and activation of the inflammatory pathways with recruitment of IFNγ-producing CD8 T cells. The inflammatory environment is associated with a decrease in production of IL15 and amplification of trophoblast injury. Loss of junctional zone trophoblasts decreases placental growth hormones and loss of IL15 contributes to a decrease in uterine NK cells. C. Downstream effects of these deficits contribute to poor placental vascularization, hypoxia, infarcts, and inflammation, culminating in fetal growth restriction and death.

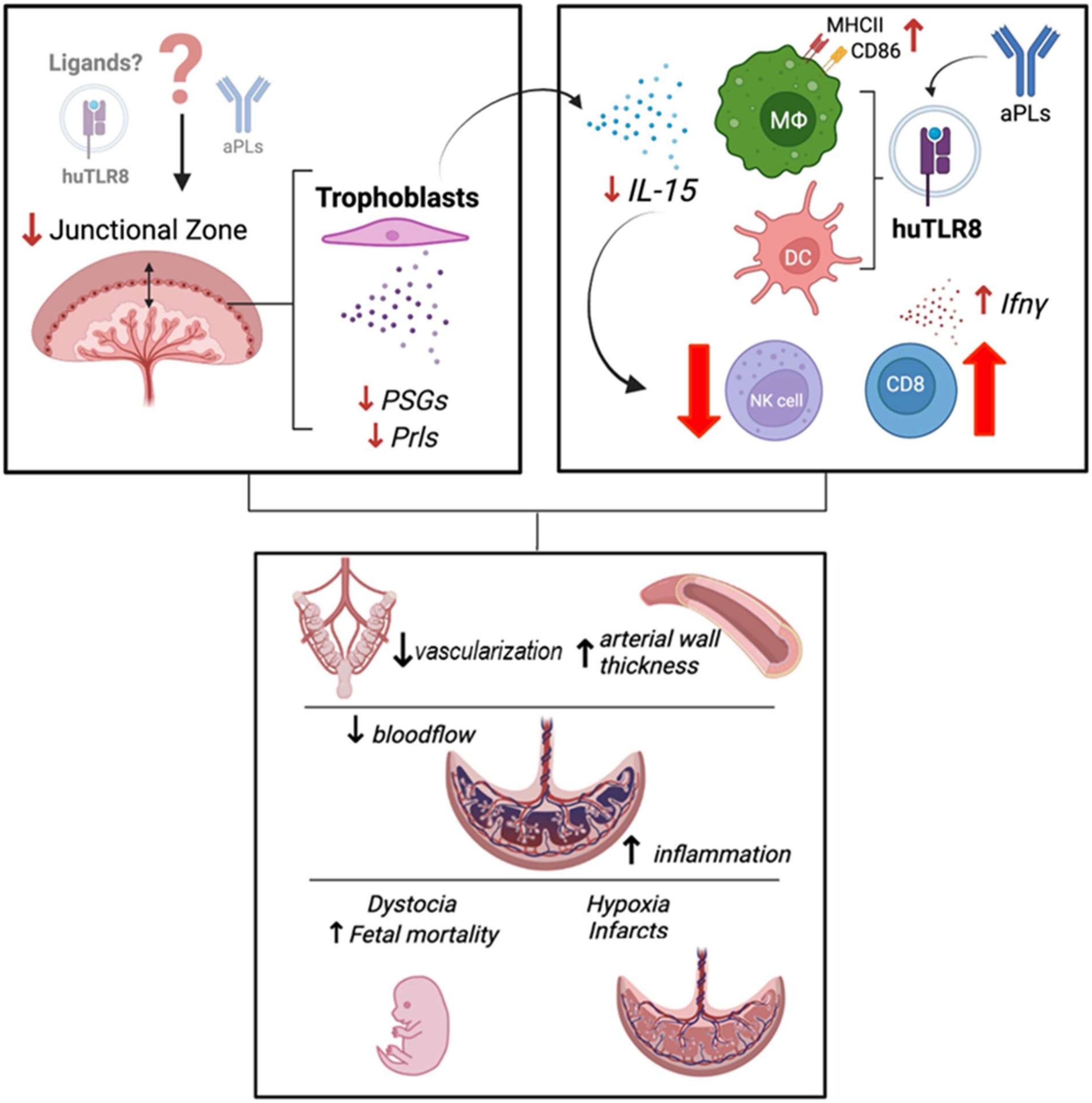

Ligands?
huTLR8
aPLs
↓ Junctional Zone
Trophoblasts
↓ PSGs
↓ Prls
MHCII
CD86
aPLs
↓ IL-15
MΦ
DC
huTLR8
↑ Ifnγ
NK cell
CD8
↓vascularization
↑ arterial wall thickness
↓ bloodflow
↑ inflammation
Dystocia
↑ Fetal mortality
Hypoxia
Infarcts
